# Canonical and novel non-canonical activities of the Holliday junction resolvase Yen1

**DOI:** 10.1101/2020.12.08.413625

**Authors:** F. Javier Aguado, Raquel Carreira, Vanesa Hurtado-Nieves, Miguel G. Blanco

**Affiliations:** Department of Biochemistry and Molecular Biology, CIMUS, Universidade de Santiago de Compostela-Instituto de Investigación Sanitaria (IDIS), Santiago de Compostela, A Coruña, 15782, Spain

## Abstract

Yen1 and GEN1 are members of the Rad2/XPG family of nucleases that were identified as the first canonical nuclear Holliday junction (HJ) resolvases in budding yeast and humans due to their ability to introduce two symmetric, coordinated incisions on opposite strands of the HJ, yielding nicked DNA products that could be readily ligated. While GEN1 has been extensively characterized *in vitro*, much less is known about the biochemistry of Yen1. Here, we have performed the first in-depth characterization of purified Yen1. We confirmed that Yen1 resembles GEN1 in many aspects, including range of substrates targeted, position of most incisions they produce or monomeric state in solution. However, we have also observed unexpected alternative processing of substrates, such as nicked HJs and a different conformational preference on intact HJs. Moreover, we demonstrate that Yen1 is endowed with additional nuclease activities, like a nick-specific 5’-3’ exonuclease or HJ arm-chopping that could apparently blur its classification as a canonical HJ resolvase. Despite this, we show that Yen1 fulfills the requirements of a canonical HJ resolvase and hypothesize that its wider array of nuclease activities might contribute to its function in the removal of persistent recombination or replication intermediates.

## INTRODUCTION

Holliday junctions (HJs) are four-way secondary DNA structures where two duplexes exchange a pair of single strands. HJs and similar cruciform structures may arise during the processes of double-strand break (DSB) repair by homologous recombination (HR), post-replication repair or during the reversal of stalled replication forks. Due to the physical linkage that HJs establish between two DNA molecules, their persistence until late stages of the cell cycle may compromise the correct segregation of sister chromatids or homologous chromosomes in both the mitotic and meiotic scenarios (1). Hence, eukaryotic cells are endowed with two main pathways to ensure their timely removal and the disengagement of the two DNA molecules. First, the combined actions of a RecQ-family helicase Sgs1 in *S. cerevisiae* (BLM in mammals), the type IA topoisomerase Top3 (TopoIIIa) and the OB-fold containing accessory protein Rmi1 (RMI1/RMI2) drive the convergent branch migration of two HJs and fuse them into a hemicatenate structure that can be subsequently decatenated. This process, double HJ dissolution, leads to the formation of non-crossover (NCO) products, exclusively (2). Second, structure-selective endonucleases (SSEs) like *S. cerevisiae* Mus81-Mms4, Slx1-Slx4 and Yen1 (MUS81-EME1/EME2, SLX1-SLX4 and GEN1) can cleave these HJs in a process generally referred to as HJ resolution, which may yield both NCOs and crossover (CO) depending on the orientation of the cuts across the HJs (3–15). Additionally, a fourth nuclease, Mlh1-Mlh3, plays a major role in the resolution of double HJs into COs to ensure chiasmata formation and accurate segregation of homologous chromosomes during meiosis I (16–19).

All these SSEs with the ability to cleave HJs are generally termed HJ resolvases. The extensive biochemical characterization of phage, prokaryotic and mitochondrial resolvases, like T4 Endo VII, T7 Endo I, *E. coli* RuvC, *S. cerevisiae* Cce1 or *S. pombe* Ydc2 defined the paradigm of “canonical” HJ resolution, whereby the enzyme introduces a pair of symmetric, coordinated nicks in opposite strands across the junction, yielding two nicked DNA duplexes that can be directly ligated without further processing (20–37). The two incisions take place within the lifetime of the protein-DNA complex, with the first one being the limiting step of the reaction and greatly accelerating the second cut (38–40). Resolvases that follow this mode of HJ resolution have been dubbed canonical resolvases. Oppositely, non-canonical resolvases typically sever HJs by non-coordinated, asymmetrical incisions that release DNA molecules containing small gaps or flaps, thus requiring additional processing before religation (41).

Yen1 and GEN1 were identified as the first eukaryotic, nuclear SSEs displaying canonical HJ resolvase activity in budding yeast and humans, respectively (10). Both belong to the subclass IV of the Rad2/XPG family of SSEs and orthologous proteins are present in most organisms analysed, with the notable exception of *S. pombe* (10, 42, 43). These enzymes are characterized by the presence of a bi-partite nuclease domain (XPG-N/XPG-I), a conserved helix-hairpin-helix motif and a recently identified DNA interaction site mediated by a chromodomain in the human and *C. thermophilum* orthologs (44–46) (Figure 1A). The biochemical analysis of various members of this subclass IV in humans, nematodes, flies, fungi and plants indicate that all these nucleases retain the characteristic 5’-flap nuclease activity of the Rad2/XPG family, but in most cases have also evolved the ability to resolve HJs in a RuvC-like manner and to process other branched DNA structures like model replication forks and nicked HJs (10, 43, 44, 46–56).

**Figure 1.**
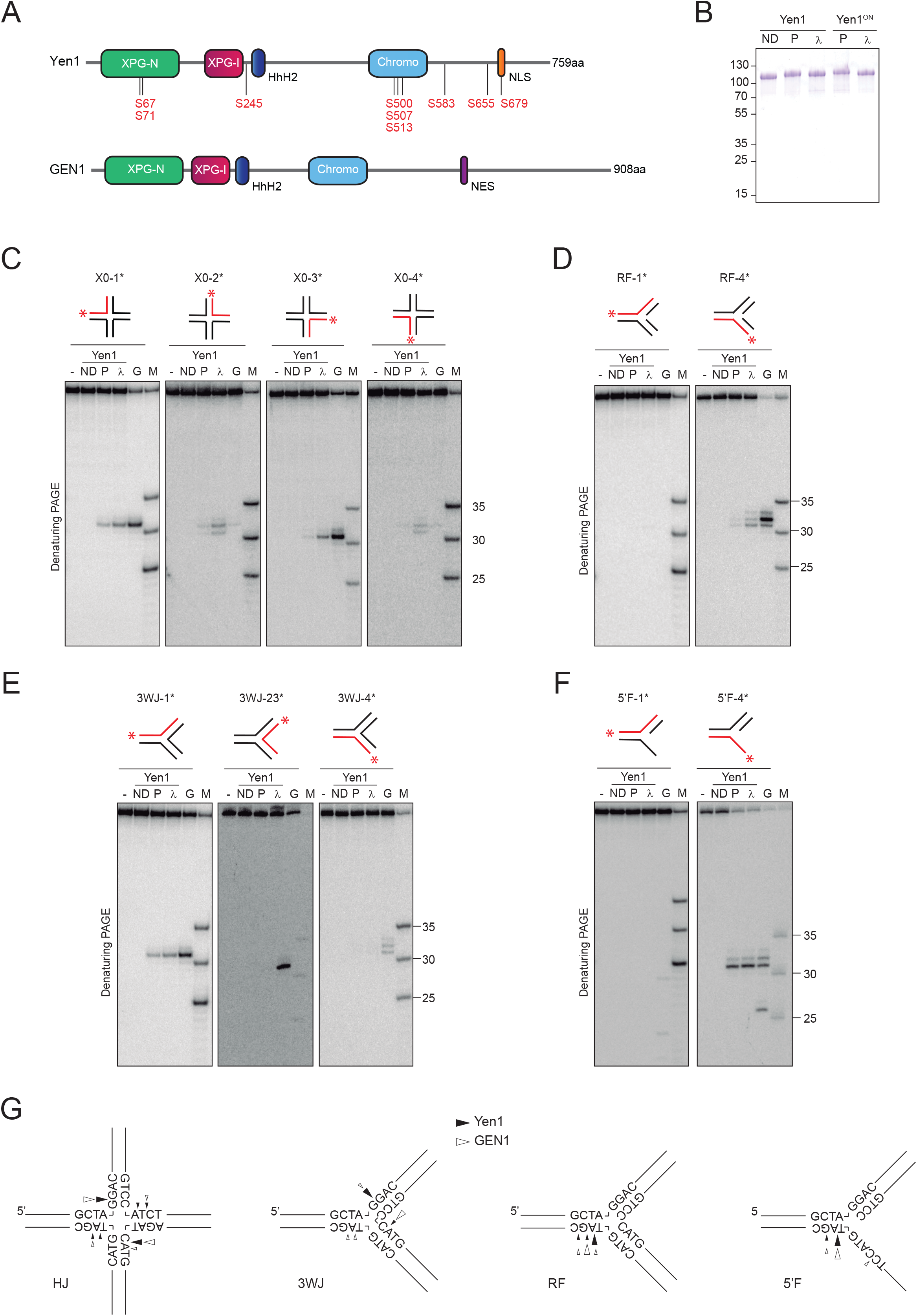
Mapping of Yen1 incisions on branched DNA substrates. (A) Graphical representation of Yen1 and GEN1 domains. Serines in consensus CDK phosphorylation sites are indicated in Yen1. (B) FTH-tagged Yen1^ND^, Yen1^P^, Yen1^λ^, Yen1^ON-P^ and Yen1^ON-λ^ were purified and analysed by SDS-PAGE and stained with Coomassie. Molecular weight markers are indicated in kDa. (C) Holliday junction X0 (20 nM) was 5’-end labelled with ^32^P (asterisk) on each strand (red) and incubated with 5 nM Yen1^ND^ (ND), Yen1^P^ (P), Yen1^λ^ (λ) or GEN1^1-527^(G) for 10 min at 30°C or 37°C, respectively. (-) indicates no enzyme. Products were analysed by 10% denaturing PAGE and phosphorimaging. A mixture of 5’-^32^P-end-labeled oligos of defined length (25, 30 and 35 nt) were used as markers (M). (D-F) As in (C), employing a (D) replication fork, (E) 3-way junction or (F) 5’-flap as a substrate. (G) Schematic representation of incision sites identified for the HJ, 3WJ, RF and 5’-flap substrates. Only the nucleotide sequence near the branch point is shown. Arrow size indicates preferential cleavage sites for Yen1 (black) and GEN1 (white).

In budding yeast, the role of Yen1 in resolution of HR intermediates has been considerably studied. While deletion of *YEN1* confers no overt DNA repair defects either on its own or in the absence of other members of the Rad2/XPG family of nucleases, it dramatically increases the DNA damage sensitivity of *mus81Δ* and *mms4Δ* mutants (57–61). Yen1 seems dispensable in meiotic recombination, but double *mus81Δ yen1Δ* display meiotic chromosome segregation abnormalities and cannot sporulate (19, 57, 62–64). These observations have consequently led to the interpretation that Yen1 acts as a back-up mechanism for Mus81-Mms4 in the removal of mitotic and meiotic recombination intermediates. However, it has been recently shown that Yen1 also has a specific role, non-redundant with Mus81-Mms4 or Slx1-Slx4, in the processing of replication intermediates that persist in the absence of the nuclease/helicase Dna2 (65, 66). Additionally, it has been shown that Yen1 subcellular localization, catalytic activity and chromatin accumulation are strictly controlled by post-translational modifications. Phosphorylation of Yen1 by the Cdc28 (CDK) kinase both reduces its DNA binding activity and promotes its export to the cytoplasm in S-phase. At the onset of anaphase, the activation of the Cdc14 phosphatase reverts this modification and allows Yen1 activation and re-ingression into the nucleus, where it acts to remove persistent recombination intermediates prior to cell division (62, 67–69). More recently, it has been observed that Yen1 can be a substrate for the SUMO ligases Siz1 and Siz2 and the SUMO-targeted ubiquitin ligase Slx5-Slx8 after DNA damage, which drive its polyubiquitination and subsequent proteasomal degradation before entry in S-phase (70). Arguably, these different layers of Yen1 regulation prevent it from targeting branched replication and recombination intermediates from S-phase until anaphase, limiting its action to persistent branched DNA intermediates in late stages of mitosis. In this sense, Yen1 mutants that bypass these regulatory mechanisms, such as Yen1^ON^ (67) (identical to Yen1^9A^ (68)) or Yen1^K714R^ (70) have proved extremely useful to ascertain the cellular consequences of premature activation or nuclear persistence of this resolvase. In particular, Yen1^ON^ has been widely employed in different experimental set-ups to uncouple the nucleolytic processing of HR intermediates from cell cycle progression in both mitotic and meiotic contexts. These studies have collectively revealed that in wild-type cells the precise timing of Yen1 activation is essential to minimize genome instability, CO formation and loss-of heterozygosity during mitosis or to properly establish CO interference during meiosis I. Contrarily, in mutants deficient in the processing of HR or replication intermediates, the sustained activation of Yen1^ON^ alleviates or suppresses the phenotypes associated to the toxic accumulation of replication or recombination intermediates (66, 67, 71–78).

Despite these advances in our understanding of Yen1 functions and regulation, our knowledge about its biochemical properties is relatively lacking compared to some of its orthologs in other organisms, as it mainly derives from its initial identification as a canonical resolvase employing whole-cell extracts (10) and a basic description of its two activation states once the protein was successfully purified (67). Here, we present the first comprehensive analysis of Yen1 biochemical properties, with a special focus in the characterization of its activities as a canonical HJ resolvase. Our results reveal that, despite its general behavior as a bona-fide canonical HJ resolvase, Yen1 is endowed with additional nucleolytic activities that may blur this adscription, at least according to its *in vitro* properties. In this sense, we have identified a novel nick-specific 5’-3’exonuclease activity that renders Yen1 resolution products unable to be ligated as well as an alternative, non-canonical two-step mode of HJ processing, which we term “arm-chopping”. Additionally, we have discovered that despite their many similarities, Yen1 and human GEN1 display different preferences for HJ strand cleavage. We speculate that such activities might contribute to the processing of particularly recalcitrant branched DNA intermediates, but also account for the detrimental effects of its premature activation.

## MATERIAL & METHODS

### Plasmids and strains

All *S. cerevisiae* strains and plasmids employed are described in Supplementary Tables S1 and S2, respectively.

The coding sequence of *YEN1*^*WT*^ in p416ADH1-YEN1^WT^-V5-6xHis (51) was mutagenized and/or subcloned into p416GPD1 (79) to create p416ADH1-YEN1^ON^-V5-6xHis, p416GPD1-YEN1^WT^-V5-6xHis and p416GPD1-YEN1^ON^-V5-6xHis. The vector pENTR221-GEN1^WT^-FTH-STOP, a kind gift from Gary Chan and Stephen C. West that contains the coding sequence of C-terminally tagged GEN1^WT^-3xFLAG-2xTEV-10xHis (FTH tag), was modified by site-directed mutagenesis and inverse PCR to generate pENTR221-GEN1^nuc^-FTH-STOP, which encodes a constitutive nuclear form of GEN1 by mutation of its nuclear export signal and addition of 3xSV40 nuclear localization signals (50). From these two vectors, derivative plasmids carrying either GEN1^WT^ or GEN1^nuc^ (pENTR221-GEN1-GO, pYES-EXP52-GEN1-V5-6xHis and pYES-EXP52-GEN1^1-527^-V5-6xHis, pAG306GAL-GEN1-FTH) were created by standard subcloning, inverse PCR and/or Gateway recombination. Details of each cloning can be provided upon request.

### Genotoxicity assays

Yeast strains were grown to mid-logarithmic phase in SC-URA containing 2% glucose. The cultures were then normalized to an OD_600_ = 0.5 and 10-fold serial dilutions were spotted onto fresh SC-URA plates containing 1% raffinose, 2% galactose and increasing concentrations of MMS. Plates were incubated in the dark for 3 days at 30°C and then photographed in a Gel Doc XR+ (Bio-Rad).

### Immunofluorescence microscopy

Strains carrying V5-tagged versions of Yen1 or GEN1, either in a WT or *mus81Δ yen1Δ* background (Supp. Table 1), were cultured in SC-URA for 16 h, then diluted to an OD_600_ = 0.2 in fresh medium and incubated for 5 hours. Cells were then fixed by addition of 0.1 vol formaldehyde and incubation at 4°C overnight. Cells were washed once in 0.1 M phosphate buffer pH 7.4, once in spheroplasting buffer (0.1 M phosphate buffer pH 7.4, 0.5 mM MgCl_2_ and 1.2 M sorbitol) and then incubated in 200 μL of spheroplasting buffer containing 28 mM β-mercaptoethanol at 30°C for 15 min. 100-T Zymolyase (AMSBIO) was added to a final concentration of 1 mg/mL and incubated at 30°C for an additional 10 min. Spheroplasts were then attached to a polylysine-coated multi-well microscope slide and dehydrated in consecutive methanol and acetone baths at -20°C. Slides were blocked for 30 min at room temperature in PBS containing 1% BSA and then incubated with mouse monoclonal anti-V5 antibodies (1:200 dilution, Invitrogen #R960-25) at 25°C for 1 h. After washing with PBS-BSA, the slides were incubated with an Alexa Fluor 488-coupled anti-mouse antibody from goat (1:200 dilution, Life technologies #A-11001). DNA was stained with DAPI. Images were acquired using a confocal microscope Leica TCS SP8 with HC PL APO CS2 63x/1.40 OIL lens, under the control of LAX software.

### Protein analysis

For the preparation of soluble protein lysates containing Yen1 or GEN1, strains expressing different versions of V5-tagged Yen1 or GEN1 were cultured in 15 mL SC-URA supplemented with 2% glucose. When a density of ∼0.6×10^7^ cells/mL was reached, cells were harvested, rinsed in water and resuspended in 15 ml SC-URA containing 1% raffinose and 2% galactose and incubated for 6 h at 30°C to induce the protein expression. Cells were then harvested and resuspended in lysis buffer (40 mM Tris-HCl pH 7.5, 150 mM NaCl, 10% glycerol, 0.10% NP-40, 1 mM DTT, 1 mM PMSF, 5 mM NaF, 5 mM H_2_NaPO_4_, 5 mM β-glycerophosphate) and disrupted using a Mini-Beadbeater-16 (BioSpec). Lysates were quantified by Bradford method, normalized to 2 mg/ml and protein expression was analysed by western blotting employing anti-V5 antibody (1:3000 dilution, Invitrogen, #R960-25).

Mobility-shift detection of phosphorylated Yen1 variants by western-blotting was performed by separating 400 ng of purified Yen1^P^, Yen1^λ^, Yen1^ON-P^ and Yen1^ON-λ^ in 7.5% SDS-PAGE gels, either unmodified or containing 40 µM Phos-tag (Wako Chemicals) and 160 µM MnCl_2_. Prior to transference, Phos-tag gels were washed with transfer buffer containing 100 mM EDTA for 30 min at RT, and then washed 3 x 15 min with transfer buffer. Proteins were detected with anti-FLAG M2-HRP (1:3000 dilution, Sigma #A8592).

### Protein purification

Wild-type Yen1, deregulated Yen1^ON^ and catalytically inactive Yen1^ND^ (E193A, E195A) were all purified as fusions with a C-terminal FTH tag, as previously described (67). Yen1^P^ represents a highly phosphorylated version of the wild-type protein, purified from cells arrested before anaphase entry in a *cdc14-1* mutant. Yen1^λ^ is a lambda-phosphatase treated version of Yen1^P^ (67). For control reactions with FTH-tagged GEN1 or GEN1^1-527^, samples for initial experiments were kindly provided by Gary Chan and Stephen C. West. Additional batches were purified as described (38). All proteins were tested for non-specific exonuclease, endonuclease, and protease contaminant activities (not shown) and analysed by SDS-PAGE through Novex Value 4-20% Tris-Glycine Mini Gel (Thermo Fisher Scientific) followed by Coomassie staining.

### Hydrodynamic analysis of Yen1

Glycerol gradient sedimentation and size exclusion chromatography were carried out essentially as described (54), with the following modifications. 25 µg Yen1^λ^ were loaded on 15%-35% glycerol gradients prepared in 40 mM Tris-HCl, pH 7.5, 500 mM NaCl, 0.1% NP-40 and 1 mM DTT. For size exclusion chromatography, 20 µg Yen1^λ^ were applied to a Superdex 200 PC 3.2/30 column (GE Healthcare) in the same buffer. In both cases, the fractions collected were analysed by SDS-PAGE through 4-12% MOPS NuPage gradient gels (Thermo Fisher Scientific) followed by Coomassie staining.

### DNA substrates

Synthetic substrates were prepared essentially as described (80), employing PAGE-purified ssDNA oligonucleotides. All the oligonucleotides are listed in Supplementary Table S3 and the strand composition of each substrate is described in Supplementary Table S4. Briefly, labelled and unlabelled oligonucleotides were mixed in a 1:3 ratio, boiled in a water bath, cooled down to room temperature overnight and fully-ligated substrates purified from 10% polyacrylamide gels. For radioactive substrates, oligos were either 5’-end labelled with [γ-^32^P]-ATP (3000 Ci/mmol, Perkin Elmer) and T4 PNK (Thermo Fisher) or 3’-end labelled with [α-^32^P]-cordycepin (3000 Ci/mmol, Perkin Elmer) and terminal transferase (New England Biolabs). For fluorescent substrates, 5’-IRDye 700, 5’-IRDye 800-labelled (Integrated DNA Technologies) or 3’-6FAM-labelled oligonucleotides (Sigma-Aldrich) were employed. Unlabelled substrates were prepared with equimolecular mixtures of oligonucleotides and purified from native 10% polyacrylamide gels by UV-shadowing. When appropriate, oligonucleotides containing 1 or 3 consecutive phosphorothioate (SP) linkages (Sigma-Aldrich) were employed.

### Nuclease assays

For the mapping of the incisions on each oligonucleotide, 5 nM protein was incubated with 20 nM unlabelled substrate (unless otherwise stated) supplemented with ∼0.1 nM 5’-^32^P-end-labeled substrate in 25 µl of reaction buffer (50 nM Tris-HCl pH 7.5, 0.5 mM MgCl_2_, 50 mM NaCl, 2% glycerol for Yen1; 50 mM Tris-HCl pH 7.5, 1 mM MgCl_2_, 25 mM NaCl, 1 mM DTT, 1% glycerol for GEN1). After 10 min of incubation at 30°C (Yen1) or 37°C (GEN1), 10 µl of each reaction were deproteinized by addition of 2.5 µl stop solution (2% SDS, 10 mg/ml proteinase K) and further incubation for 20 min at 37°C. Another aliquot of 10 µl was mixed with 1 volume of 2X denaturing loading buffer (TBE 1X, 80% formamide) and incubated at 99°C for 3 min. The radiolabelled products were then separated by PAGE through 10% native or denaturing (7M urea) gels. Radiolabelled molecules were analysed by phosphorimaging using a Typhoon FLA 9500 and quantified by densitometry using ImageQuant software (GE Healthcare). In experiments employing an X0 HJ with one SP in strand X0-3 (X0-SP), 5 nM unlabelled substrate was used, and the incubation time was 5 min for Yen1^λ^ and 2.5 min for GEN1. In experiments employing an X0 HJ with 3 SP in strands X0-2 and X0-4 (HJ-2SP) 5 nM unlabelled substrate was used and 30 min incubation for Yen1 and 2.5 min for GEN1.

To assess the preference for the axis of HJ resolution, reactions were carried out as described for the mapping analyses. 5 nM unlabelled J3 HJ spiked with ∼0.5 nM 5’-^32^P-end-labeled junction was incubated with 5 nM Yen1^λ^ (30°C, 10 min) or GEN1 (37°C, 5 min). The radiolabelled products were separated by 12% denaturing PAGE and quantified by phosphorimaging as described.

For religation experiments, ∼0.5 nM 5’-^32^P-end-labeled asymmetric HJ X1-T (38) or X1-T-SP were incubated with 10 nM Yen1 or GEN1 for 30 min under the same conditions as described above. 10 µl aliquots of each reaction were either left untreated or adjusted to 1X T4 ligase buffer, supplemented with 5U of T4 DNA ligase (Fisher, #EL0014) and further incubated for 1 h at 25°C. All reactions were stopped by adding 1 vol of 2X denaturing loading buffer and incubated at 99°C for 3 min. Products were then analysed by 10% denaturing PAGE and phosphorimaging as described above.

For exonuclease assays, ∼0.5 nM of either 5’- or 3’-^32^P-end-labeled nicked or gapped dsDNA substrates were incubated with increasing concentrations of Yen1 (0, 0.25, 0.5, 1, 2, 4, 8 nM in Figure 5C; 0, 0.5, 2, 8 nM in Figure 5D) for 10 min at 30°C in a total volume of 20 µl. After incubation, all reactions were stopped with 5 µl stop buffer, 37°C, 1 h. For analysis under native conditions, reactions were mixed with 0.2 vol 6X native loading buffer (15% Ficoll-400, 60 mM EDTA, 19.8 mM Tris-HCl pH 8.0, 0.48% SDS), and separated in 10% PAGE native gels. For analysis under denaturing conditions, reactions were mixed with 1 vol of 2X denaturing loading buffer, boiled and loaded on12% PAGE gels containing 7M urea.

Exonuclease experiments with 6FAM-fluorescent substrates (Figure 5E) were carried out with 50 nM enzyme, 10 nM substrate and incubated at 30°C for Yen1 or 37°C for GEN1 for 10 min. Reaction products were separated through 13% PAGE gels containing 7M urea and visualized in a Typhoon FLA 9500.

Time-course experiments for Yen1 were carried out using 50 nM IRDye-800 fluorescent DNA substrates and 40 nM Yen1 for reactions with X0 HJ or RF substrates and 10 nM Yen1 in those with the 5’-flap substrate. Reactions were incubated at 30°C, aliquots taken at each time point and stopped by addition of 2.5 µl stop solution and further incubation at 37°C for 30 min. Products were separated through 10% native PAGE and analysed using an Odyssey infrared imaging system and Application software (LI-COR Biosciences).

Nuclease assays with whole-cell lysates from strains expressing Yen1 or GEN1 were performed with IRDye-700 labelled X0 as stated above. 1 µl of normalized lysate (∼2 mg/ml total protein) was added to each reaction and incubated for 1 h at 30°C. Products were analysed by 10% native PAGE and the gels were scanned using an Odyssey infrared imaging system (LI-COR Biosciences).

### Cruciform assays

In experiments with the cruciform-forming pIR9 plasmid (48), extrusion of the cruciform was favored by preincubation at 37°C in extrusion buffer (50 mM Tris-HCl pH 7.5, 50 mM NaCl, 0.1 mM EDTA). Cruciform cleavage assays were carried out in 10 µl Yen1 reaction buffer containing 4.5 nM plasmid and 90 nM of Yen1 or 50 nM GEN1. Resistance to digestion by *Eco*RI was employed to measure cruciform extrusion levels. Products were separated in 0.8% agarose gels, stained with ethidium bromide, and imaged and quantified on a Gel Doc XR + System (Bio-Rad). For the labeling experiments, reactions were carried in a final volume of 20 µl, from which half of the reaction was loaded on agarose gels as described, and the other half was denatured at 99°C for 3 min. After this, 50 mM Tris-HCl was added to the reaction for a final volume of 60 µl to pass through a G-25 MicroSpin column. Following this step, reaction products were labeled by adding 1 µl [γ-^32^P]-ATP and T4 PNK for 1 h at 37°C. The non-incorporated isotope was eliminated using G-25 columns and the eluted DNA was precipitated with 2 vol of ethanol, 0.1 vol of 3 M sodium acetate pH 5.2 and 0.02 vol of 5 mg/ml glycogen. The pellet was resuspended in 10 µL 2X denaturing loading buffer and samples analysed in 15% denaturing gels as described above.

For the ligation experiment, pIR9 was incubated with Yen1 as previously described. In addition, incubation of the cruciform plasmid with Nt.BspQI (NEB) was used as a control for a nicked circle. After the nuclease reaction, the DNA was purified by using commercial DNA purification kits (E.Z.N.A.Cycle-Pure Kit, Omega Bio-tek #D6492-02). Following the purification, 5 U T4 DNA ligase were added to 100 ng of purified DNA in the supplemented buffer, for 1 h at 37°C. Products were analysed as stated above.

## RESULTS

### Yen1 and GEN1 cleave four- and three-way junctions at similar, but not identical, positions

Yen1 nuclease activity is regulated by CDK-dependent phosphorylation events that inhibit both its overall DNA-binding and catalytic activity (67, 68). However, it is not known whether these changes may also alter the position of the incisions that Yen1 creates on the substrates. To investigate this, we purified highly phosphorylated (Yen1^P^) and *in vitro*, lambda phosphatase-dephosphorylated (Yen1^λ^) versions of Yen1 (Figure 1B) and mapped the cleavage sites they introduced on several branched DNA structures, including HJs, 3-way junctions (3WJ), replication forks and 5’-flap substrates employing 7M urea denaturing PAGE (Figure 1C-F). The same reactions were also analysed by native PAGE (Supp. Fig. 1). We observed that, despite their different efficiency, the incision sites produced by Yen1^P^ and Yen1^λ^ on all the tested substrates are entirely coincident, indicating that the upregulation of Yen1 activity by dephosphorylation does not alter its cleavage specificity.

In parallel, we also compared Yen1 incisions with those created by its human homolog GEN1 on the same set of substrates (Figure 1C-G and Supp. Fig. 1). On the static, fully-ligated X0 HJ substrate (10), GEN1 typically introduces symmetrical nicks 1 nt at the 3’of the branching point specifically on strands 1 and 3 (Figures 1C and 1G), in agreement with previous reports (10, 38, 54). On top of these cuts, Yen1 can nick strands 2 and 4 of X0, typically 1 or 2 nt at the 3’ of the branching point. Therefore, it seems that while GEN1 preferentially resolves the X0 HJ over the 1-3 axis, Yen1 can process it across both the 1-3 and 2-4 axis. Interestingly, the analysis of the cleavage products by native PAGE revealed that Yen1, but not GEN1, gives rise to two additional species apart from the nicked duplex expected by canonical resolution (Supp. Fig. 1). These species migrate in positions comparable to those of a replication fork-like (RF-like) structure and a 30 nt duplex, which would be consistent with the cleavage of two annealing strands, and thus releasing one helical arm of the junction. Differences between the two enzymes in their preference to cleave specific strands can also be observed on a fully continuous 3WJ (Figures 1E and 1G). For this substrate, Yen1 has a marked preference to incise strand 1, again 1 nt at the 3’ of the branching point, while GEN1 produces detectable nicking on all strands of this substrate. Together with the results obtained with the X0 HJ, this suggests that the two enzymes may recognize or remodel these continuous branched substrates in a different manner. However, substrates with discontinuous strands like model replication forks (RFs) or 5’-flaps were almost exclusively processed by both Yen1 and GEN1 on strand 4, 1 or 2 nt at the 3’ from the branching point of the 5’ arm/flap (Figures 1D, 1F and 1G). An additional incision 4 nt into the 5’-flap strand for GEN1 that has been previously observed was also recapitulated in our analysis (Chan and West, 2015). Additionally, we also tested if Yen1 could nick any of the strands of 3’-flap, splayed arm, double- and single-stranded DNA substrates. In agreement with our previous results showing no cleavage of these species (67), we did not detect incisions produced by Yen1 on any of their constituent strands (Supp. Fig. 2).

### Yen1 can incise both the opposing and 5’ strands of nicked HJs

Nicked HJs are important intermediates during HR, being the precursors of fully ligated HJs and regarded in many instances as the key intermediate to generate crossovers (81, 82). To specifically assess how Yen1 resolves these structures, we generated the 4 possible nicked HJs based on the X0 structure (X0n1 to X0n4, with 1-4 indicating the nicked strand) and independently labelled each of the intact strands. The resulting set of 12 substrates was incubated with Yen1 or GEN1 and their incisions mapped as described in the previous section. When we labelled the strand opposing the nick (Figure 2A), we observed that both Yen1 and GEN1 hydrolysed it 1 nt to the 3’-side of the junction point (position 31) for X0n3. In junctions X0n1, X0n2 and X0n4, Yen1 still cleaves mainly at position 31, but GEN1 does it 1-3 nt to the 3’-side of the junction point, mainly in position 32 of the oligo (Figures 2A and G). Interestingly, these incisions on the nicked X0 are not perfectly symmetrical to the pre-existing nick in position 30 and thus would lead to duplex products that would contain typically 1 nt (up to 3 nt) gaps or 3’-flaps, as it has been observed for the *Arabidopsis* orthologs (48).

**Figure 2.**
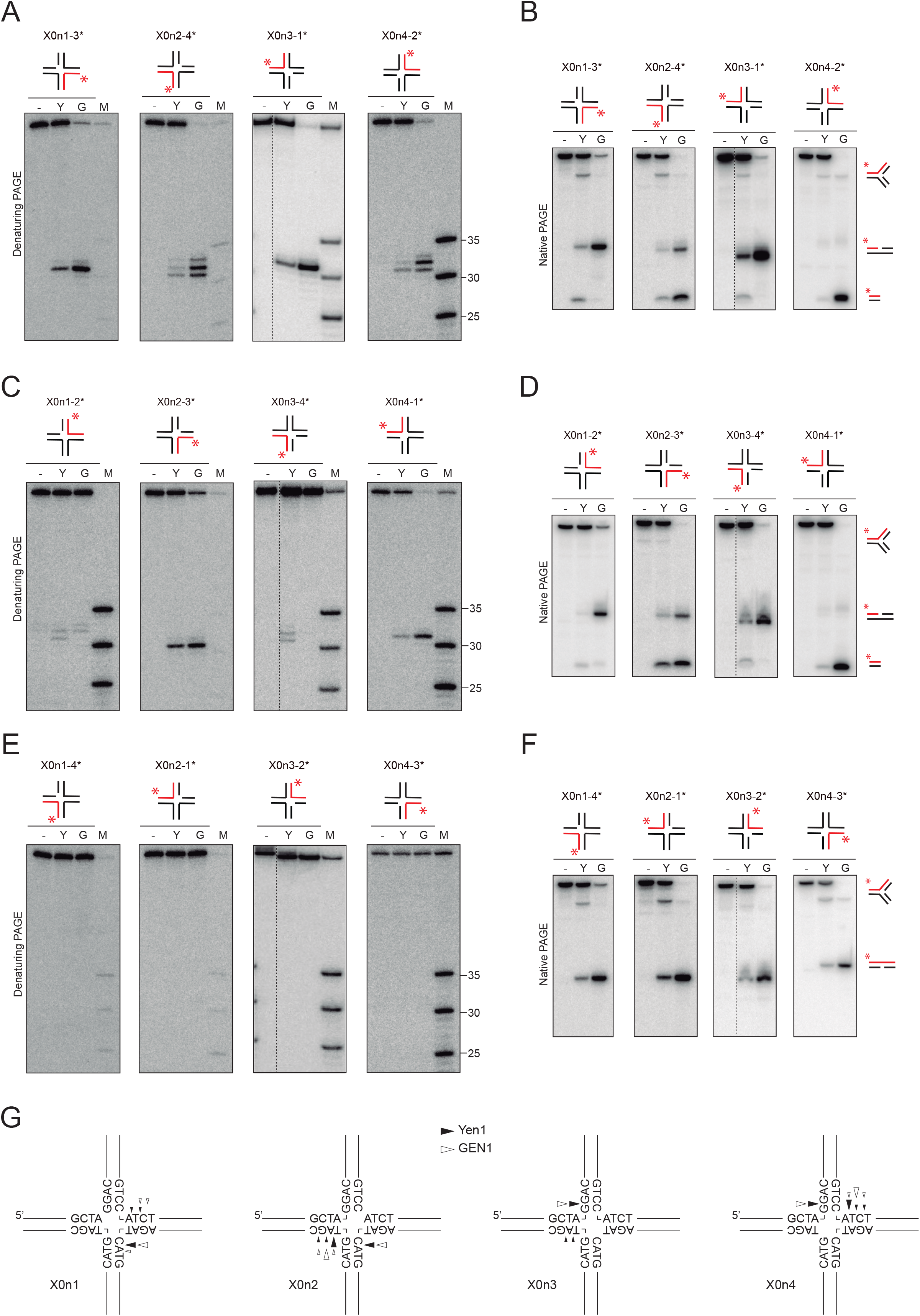
Mapping of Yen1 incisions on nicked HJs. Reactions with different nicked X0 HJs and Yen1^λ^ or GEN1^1-527^ were carried out and labelled as in Figure 1. (A) Reactions using nicked X0 substrates labelled on the strand opposite the nicked were analyed by 10% denaturing PAGE or (B) 10% native PAGE (cleavage products depicted on the right). (C-D) same as (A-B), but employing nicked X0 substrates labelled on the 5’strand with respect to the nick. (E-F) same as (A-B), but employing nicked X0 substrates labelled on the 3’strand with respect to the nick. Dotted lines in some gels indicate splicing of superfluous lanes. (G) Schematic representation of incision sites identified for the different nicked X0 substrates. Please note the discontinuity of one strand in each of them. Only the nucleotide sequence near the branch point is shown. Arrow size indicates preferential cleavage sites for Yen1 (black) and GEN1 (white).

It is generally considered that the presence of a nick in a HJ increases its flexibility and the accessibility of the SSEs for the cleavage of the opposite strand, thus favouring counter-nicking and resolution of the junction (83). Accordingly, analysis of the same reaction products on native gels also revealed that the expected 60 nt duplex resulting from resolution of the X0n1 and X0n3 was almost the exclusive product for GEN1 (Figure 2B). However, in the case of Yen1, the same pattern of 3 bands (RF-like, nicked duplex, small duplex) observed with intact X0 was still present, to different extents, with all the nicked X0 versions. Surprisingly, the major product of GEN1 on X0n2 and X0n4 was the small duplex molecule, which necessarily involves the cleavage of the 5’strand with respect to the nick. Indeed, when we labelled the 5’strand in all the nicked HJs, we detected mild cleavage by both nucleases in all cases, except for GEN1 on X0n3-4* (Figure 2C and 2G) and consistently, we observed again the presence of the small duplexes in the native gels, except for X0n3-4* (Figure 2D). Conversely, we detected no incisions or the presence of the small duplex product when the nicked junctions were labelled on the 3’strand with respect to the nick (Figures 2E and 2F).

The removal of one arm from a nicked HJ to produce a three-arm structure has already been observed for the *A. thaliana* orthologs of Yen1, AtGEN1 and AtSEND1 (48), being referred to as *Ref-I* (Replication Fork Intermediate) activity. Interestingly, the small duplex that we observe when we label the strand opposite the nick (Figure 2B) cannot arise directly as a consequence of this *Ref-I* activity. Instead, it must involve a sequential reaction, in which the first incision by *Ref-I* activity on the 5’strand with respect to the nick is followed by a second incision on the resulting RF-like intermediate, with the typical 5’ polarity of the Rad2/XPG family (Supp. Fig. 3A).

### Both symmetrical resolution and arm-chopping contribute to the formation of nicked duplex DNA from a synthetic HJ

The paradigm of canonical HJ resolution proposes that two symmetrical, coordinated incisions across the axis of the HJ produce a pair of nicked duplexes (21, 84). Surprisingly, when we analysed the cleavage products generated by Yen1 on an intact X0 HJ in native gels (Supp. Fig. 1), we detected the formation of, apparently, the same two additional species that we observed with the nicked X0 HJs: one with a migration compatible with a RF-like structure and another migrating like a small (∼30 nt) DNA duplex.

This suggests that Yen1 may exhibit an alternative way of processing an intact HJ: the cleavage of both strands in one arm of the HJ, releasing a small dsDNA fragment and a RF-like, three-arm structure (Supp. Fig. 3B). This arm-chopping activity has also been observed for some of Yen1 orthologs such as OsGEN1 (56), AtGEN1 and AtSEND1 (48) and DmGEN1 (49). Importantly, the RF-like structure produced by this arm-chopping would be susceptible of further processing to yield a nicked dsDNA product, similar to the one expected by symmetrical resolution, in a situation analogous to that described for the *Ref-I* activity on nicked HJs (Supp. Fig. 3A). This poses an interesting caveat: is Yen1 able to symmetrically resolve an intact HJ into two nicked dsDNA molecules or do these nicked duplexes arise from a two-step mechanism involving arm-chopping? To address this question, we generated an X0 HJ variant in which the opposing oligos X0-2 and X0-4 were modified to contain three hydrolysis-resistant phosphorothioate linkages (SP) between nucleotides 30, 31, 32 and 33 (Figure 3A). Therefore, this structure could only be processed by symmetrical resolution, i.e. by the introduction of two nicks on oligos X0-1 and X0-3, respectively. After confirming that oligos X0-2 and X0-4 where resistant to cleavage by both Yen1 and GEN1 (Figure 3B), we compared the ability of both resolvases to process this structure with respect to an unmodified X0 HJ and verified that the ability of each enzyme to introduce a nick at nucleotide 31 in the oligo 1 was similar for both substrates (Figure 3C). Remarkably, when analysing the same reactions on native gels, we observed that both the RF-like and small duplex products that Yen1 generates on the X0 disappear when the X0-2SP is employed, while most of the nicked duplex product remained (Figure 3D). We quantified the formation of nicked duplex and total cleavage (RF-like + nicked duplex + small duplex) in each reaction and observed that, for GEN1, total cleavage is equivalent to nicked duplex formation, as expected for a canonical resolvase. For Yen1, the nicked duplex represents approximately 2/3 of the total cleavage (19% vs 28%) on a normal X0, with an undetermined contribution from each mechanism. However, on the X0-2SP virtually all cleavage products correspond to the nicked duplex (14%), which in this case can only arise from symmetrical resolution (Figure 3D). Therefore, these results demonstrate that Yen1 is capable of symmetrical resolution of an intact HJ, despite being endowed with an alternative arm-chopping mode of HJ processing.

**Figure 3.**
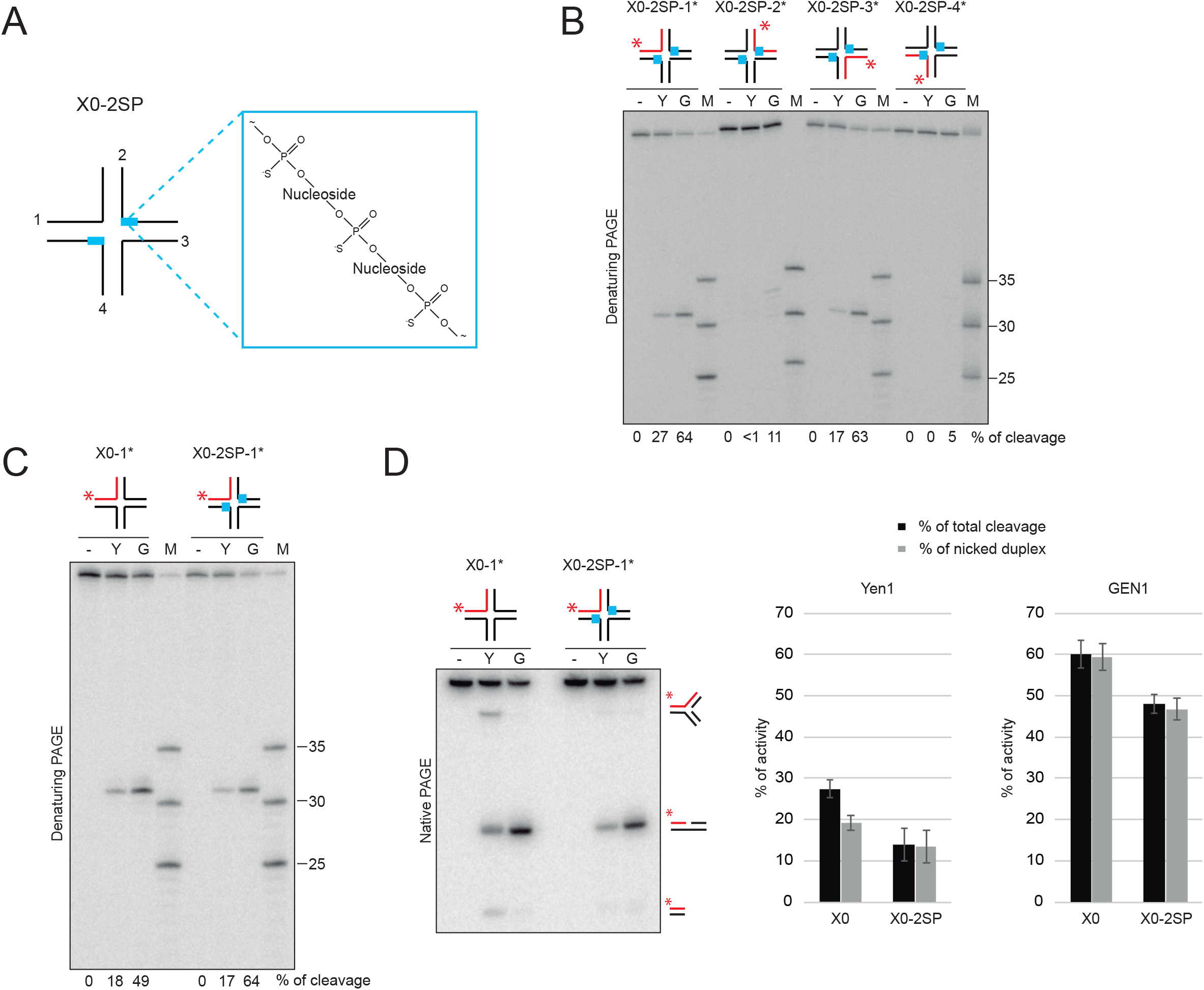
Yen1 can resolve HJs symmetrically. (A) Scheme of the SP-containing X0 HJ employed in the assay (X0-2SP), which contains 3 hydrolysis-resistant phosphorothioate linkages in positions 30, 31, 32 and 33 of oligos 2 and 4 (indicated by a blue box). (B) Reactions were carried out and labelled essentially as in Figure 1, using 5 nM X0-2SP, 5’-^32^P -labelled (asterisk) and 5nM Yen1^λ^ (10 min, 30°C) or GEN1^1-527^ (5 min, 37°C). Reaction products were analysed by 10% denaturing-PAGE followed by phosphorimaging and quantified using ImageQuant. (C) Reactions with X0 or X0-2SP (5 nM) 5’-^32^P -labelled in oligo 1 were incubated with Yen1^λ^ or GEN1 in the same conditions as in (B) and analysed by 10% denaturing PAGE. (D) The same reactions as in (C) were analysed by 10% native PAGE (left panel). The formation of nicked dsDNA products and the total amount of all cleavage products were quantified (right panel). Data represented as mean values ± SD (n = 3).

### Yen1 and GEN1 display a different preference in strand cleavage on a HJ with a fixed conformation

In the presence of Mg^2+^, HJs undergo a conformational change from an open, square structure into a form where pairs of arms are coaxially stacked (85). This implies the existence of two DNA strands that will appear as continuous as opposed to two exchanging strands. Some HJ resolvases like RuvC, T7 Endo I and GEN1 show a preference for the cleavage of the continuous strands (32, 54, 86, 87), while Hjc or T4 Endo VII incise the exchanging strands (88–90). To assess if Yen1 displays any bias in cleavage orientation, we determined the incision sites it produced on a J3 synthetic HJ. In solution, this junction exists almost exclusively as a conformer with arm-stacking of 2 over 4 and 1 over 3 (Figure 4A) (31). As previously reported, GEN1 preferentially nicks the *x* and *h* strands (Figure 4B-D) (54). Unexpectedly, and despite a comparatively low activity on this substrate, Yen1 processed the J3 HJ through incisions in the *b* and *r* strands (Figure 4B-D). Therefore, these results indicate that, like GEN1 and other resolvases, Yen1 displays a marked preference bias for the cleavage of one particular axis of this J3, which could be due to a different docking mode on the junction or sequence preference compared to its human counterpart.

**Figure 4.**
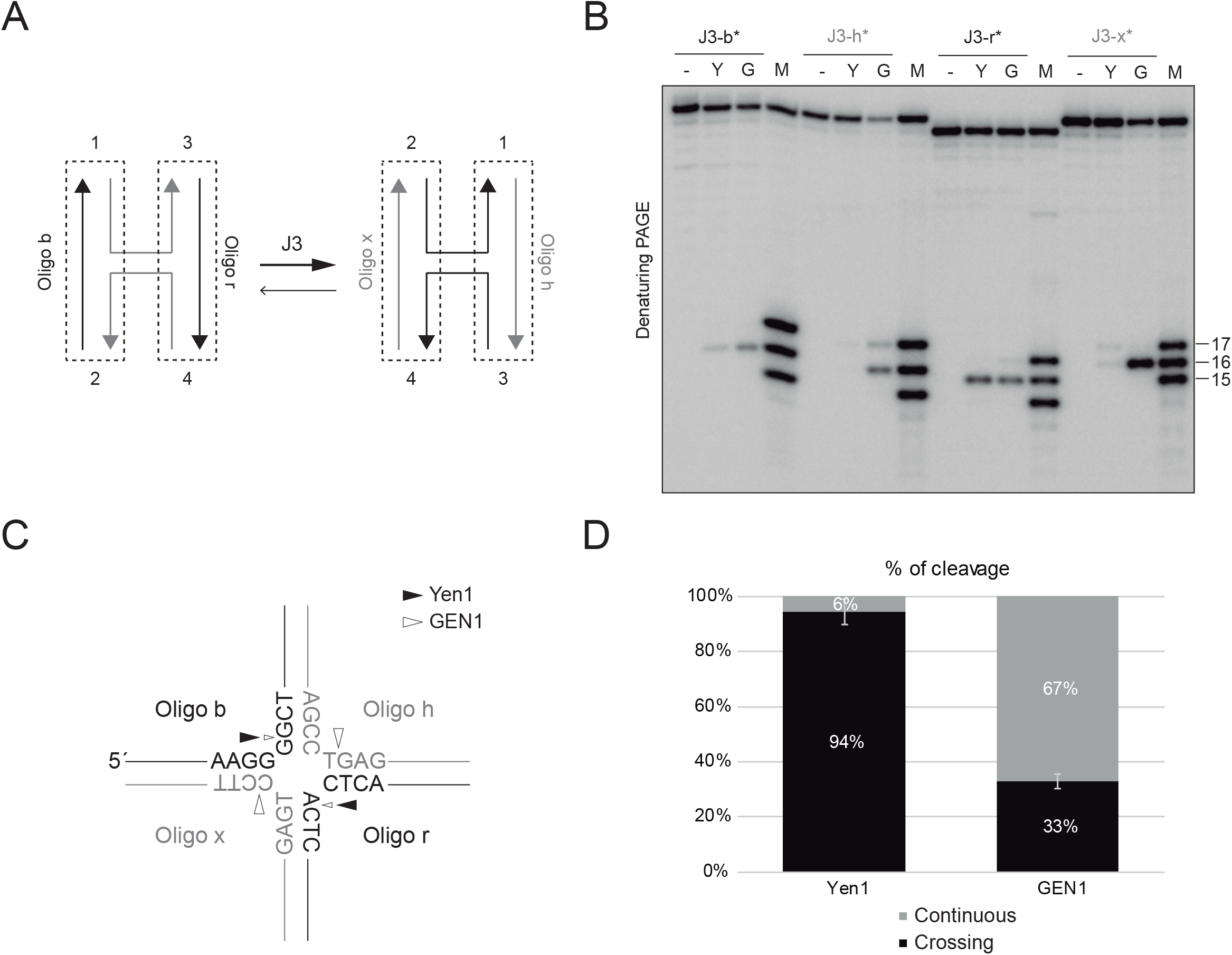
Yen1 shows a bias for the cleavage of the crossing strands of a HJ. (A) Representation of the stacking conformers of the J3 HJ substrate. In its predominant state, oligos x and h (grey) behave as continuous strands while b and r (black) are crossing strands. (B) 5 nM J3-HJ 5’-^32^P -labelled on the indicated strands was incubated with 5 nM Yen1^λ^ (30°C, 10 min) or GEN1^1-527^ (37°C, 5 min). (-) control reaction without enzyme. Reaction products were analysed by 12% denaturing PAGE and phosphorimaging. A mixture of 5’-^32^P-end-labelled oligos of defined length (15, 16 and 17 nt) were used as markers (M). (C) Schematic representation of the main cleavage sites on the J3 HJ by Yen1 (black arrows) and GEN1 (white arrows). (D) Quantification of the proportion of the cleavage of crossing vs continuous strands from (B). Data represented as mean values ± SD (n = 3).

### A novel 5’-3’ exonuclease activity in Yen1 prevents the ligation of its HJ resolution products

GEN1 was first characterized as a “canonical” HJ resolvase due to its ability to introduce symmetrical nicks that lead to the formation of nicked dsDNA molecules that can be ligated without any further processing (10). As we have shown, Yen1 can also introduce symmetrical incisions, but its ability to produce resolution products that can be directly ligated remains unaddressed. Therefore, we decided to test if HJ resolution by Yen1 also fulfils this condition. For this purpose, we employed the X1-T asymmetrical HJ that allows the ligation of cleavage products created by GEN1 (38). Resolution of X1-T across strands 1 (53 nt, 5’-labelled) and 3 (60 nt) and further religation of the nicked product with T4 DNA ligase creates a new hybrid 5’-labelled strand of 60 nt that can be detected by denaturing PAGE (Figure 5A). Using this set-up, we confirmed that GEN1 resolution products can be readily ligated (>40%) (Figure 5B), as previously reported (10, 38, 54). However, ligation products were almost undetectable for Yen1. Given that the nicks produced on this junction are symmetrical and thus *a priori* able to be sealed, we speculated that Yen1 might be further processing its cleavage products. To test this hypothesis, we created nicked dsDNA substrates and determined whether Yen1 could act on them. When we labelled the 5’-phosphate group at the nick, we observed that incubation with Yen1 resulted in the loss of the labelling, as determined by the appearance of a fast migrating band in both native and denaturing gels (Figure 5C, left panel). Conversely, when labelling the 3’-end of the same strand downstream the nick, Yen1 produced a slight shift of the substrate in native PAGE and the degradation of the substrate up to 27 nucleotides in denaturing PAGE (Figure 5C, right panel). Both results indicate that Yen1 is endowed with a 5’-3’ exonuclease activity specific for nicked DNA molecules. To further characterize this previously undescribed exonuclease activity, we created a series of 3’-labelled dsDNA substrates with gaps of increasing length and checked if Yen1 could extend such gaps (Figure 5D). We observed that nicks and gaps up to 2 nt could be efficiently extended into 3 nt gaps. However, 3 nt gaps became poor substrates for Yen1 exonuclease activity and 4 or 5 nt gaps were not further extended, indicating that, in this context, it acts as a low processivity exonuclease. Consistently with the exonuclease nature of this activity, addition of an SP linkage between the first two nucleotides of the downstream oligo in a nicked dsDNA, prevented Yen1 from processing this substrate (Figure 5E). Interestingly, the same experiment also revealed that GEN1 is endowed with the same activity, albeit it is comparatively weak, excising only 1 nt. Finally, we hypothesized that if this exonuclease activity was processing the resolution products of Yen1 on the X1-T HJ and preventing their ligation, the inclusion of an SP linkage between nt 31-32 in the oligo 3 of this X1-T HJ (1 nt at the 3’of the incision for resolution) should prevent this activity and allow their ligation. Indeed, when the resolution products created by Yen1 on such X1-T-SP were incubated with T4 ligase, the 60 nt ligated strand could be detected, unlike with the unmodified X1-T HJ (Figure 5F). Altogether, these results reveal the existence of a 5’-3’ exonuclease activity in Yen1 that specifically extends nicks in dsDNA molecules into single-stranded gaps up to 3 nt long. Such activity efficiently processes the products of symmetrical resolution by Yen1, rendering them unable to be ligated *in vitro*.

**Figure 5.**
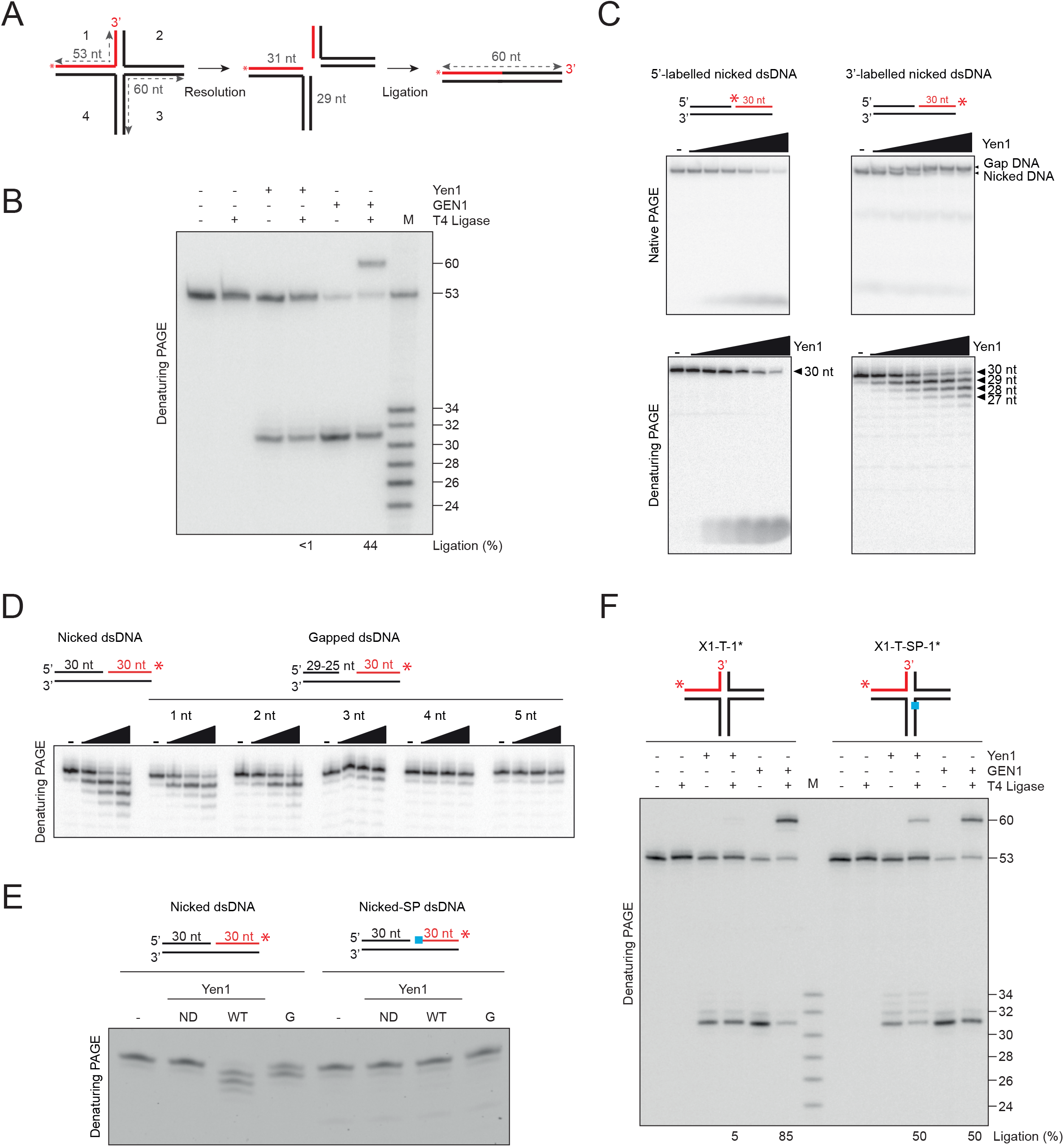
A 5’-3’ nick-specific exonuclease activity prevents the religation of the resolution products by Yen1. (A) Scheme of the religation experiment employing the asymmetrical X1-T HJ substrate. Symmetric cleavage of this substrate, which contains a 53 nt, 5’-labelled oligo (red), and its subsequent ligation generates a new 60-nt labelled strand. (B) 1 nM X1-T was incubated with 10 nM Yen1 or GEN1 for 30 min at 30 °C or 37°C, respectively. Reactions were then supplemented with ligation buffer and T4 DNA ligase (5 U) and incubated for 1 h at RT. Products were analysed by 10% denaturing PAGE followed by phosphorimaging and quantified using ImageQuant. A mixture of 5’-^32^P-end-labeled oligos of defined length (53, 34, 32, 30, 28, 26 and 24 nt) were used as markers (M). (C) 0.5 nM nicked dsDNA, either 5’-^32^P-end (left panels) or 3’-^32^P-end (right panels) labelled were incubated with increasing concentrations of Yen1^λ^ (0, 0.25, 0.5, 1, 2, 4, 8 nM) for 10 min at 30°C. Products were then analysed by 10% native (top panels) or denaturing (bottom panels) PAGE and phosphorimaging. (D) 0.5 nM of 3’-^32^P-labelled dsDNA substrates with gaps of increasing length were incubated with increasing concentrations of Yen1 (0, 0.5, 2, 8 nM) for 10 min at 30°C. Products were analysed as in (C). (E) 50 nM nicked dsDNA or exonuclease-resistant (SP-linkage in the 5’ nucleotide, in blue) nicked dsDNA, 3’-6FAM-end labelled on the indicated strand (red) were incubated with Yen1^ND^, Yen1^λ^ and GEN1 (10 nM) for 10 min at 30°C or 37°C, respectively. (-) indicates no enzyme. Fluorescent products were analysed by 13% denaturing-PAGE and scanned at 488 nm in a Typhoon FLA 9500. (F) As in (B), but employing both the unmodified X1-T HJ substrate and a SP-containing variant in position 31 of its oligo 3.

### Yen1 is a monomer in solution

Coordinated HJ resolution requires the introduction of near-simultaneous nicks across one of the axes of the structure. In bacteria and archaea, this is facilitated by the dimeric nature of HJ resolvases in solution (41). However, Yen1 orthologs are monomers in solution, like all the members of the Rad2/XPG family and, hence, it has been proposed that coordinated incisions require their dimerization on the HJs (38, 49, 51, 54). To assess the state of Yen1 in solution, we analysed its hydrodynamic properties by size exclusion chromatography and sedimentation through glycerol gradients (Supp. Fig. 4). The observed values of Stokes radius (4.9 nm) and S value (3.9) for Yen1, allowed us to estimate an apparent molecular weight of 80 kDa in solution, which would be consistent with a monomeric state, as reported for other orthologs.

### Yen1 can process a plasmid with a cruciform structure by both resolution and arm-chopping

Canonical HJ resolution involves the introduction of a pair of nicks in a coordinated manner, this is, within the lifetime of the enzyme dimer-HJ complex. The ability of numerous resolvases to introduce coordinated cuts has been extensively tested employing plasmid-based substrates that extrude palindromic sequences as cruciform, like pIRbke8 derivatives (54, 91) or pIR9 (48). Coordinated cleavage of the cruciform structure leads to the linearization of the plasmid, while individual nicks on the cruciform allow the release of negative supercoils and result in the reabsorption of the cruciform into a relaxed nicked circular DNA (Figure 6A). It has been shown that dimers of human GEN1 and CtGEN1 introduce coordinated nicks on the cruciform within the lifetime of the enzyme-DNA complex, leading almost exclusively to the formation of linear products (38, 51, 54). We wanted to address if Yen1 also displays a similar level of coordination when processing this substrate. Interestingly, we observed that the incubation of the cruciform-containing plasmid pIR9 with Yen1 gave rise to a mix of products depending on the phosphorylation status of the enzyme: the phosphorylated version of Yen1 resulted mostly in its conversion to the slowest migrating band, the relaxed circular form of the plasmid (Figure 6B), while the major product after incubation with the more active, dephosphorylated version of Yen1 was the linearized vector. These results suggest that the activation of Yen1, in addition to the enhancement of its nuclease activity, may also improve its ability to carry out coordinated cleavage within the lifetime of the enzyme dimer-DNA complex.

**Figure 6.**
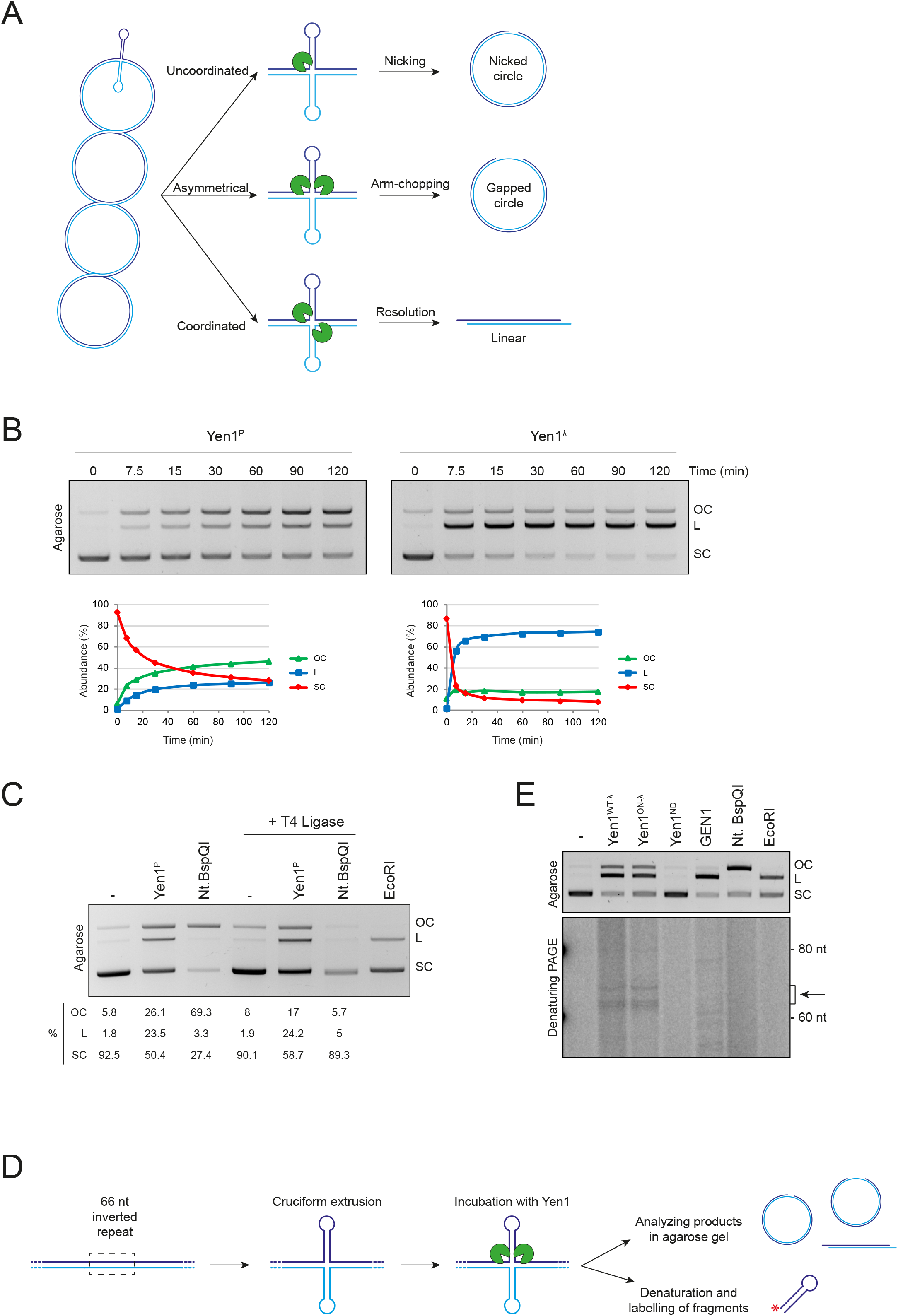
Yen1 processes plasmid-borne cruciform structures by coordinated incisions. (A) Schematic representation of the cruciform cleavage assay used to determine whether incisions are coordinated or not. A third possibility incorporating arm-chopping (central pathway) is added to the two classical outcomes of the assay. (B) 4.5 nM pIR9 plasmid was incubated with 90 nM Yen1^P^ or Yen1^λ^ at 30°C for the indicated times. Products were analysed in 0.8% agarose gels and visualised by EtBr staining in a Gel Doc Xr+ (Bio-Rad). Quantification was performed by using the Image Lab software (Bio-Rad). OC: open circle; L: linear; SC: supercoiled. (C) Ligation experiment with pIR9. 4.5 nM pIR9 was incubated with 90 nM of Yen1^P^ or 10 U Nt.BspQI for 1 h at 30°C and 50°C, respectively. Products were then purified, treated with T4 ligase for 1 h at 37°C, and then analysed and quantified as in (B). EcoRI was used as a control of the extrusion level of the plasmid. (D) Scheme of the experimental set-up to detect the hairpins released by arm-chopping on pIR9. (E) 4.5 nM pIR9 was incubated with 90 nM of different versions of Yen1 (Yen1^λ^, Yen1 ^ON-λ^ and Yen1^ND^), 50 nM of GEN1, 10 U of Nt.BspQI or EcoRI for 90 min. Half of the reaction was loaded on a 0.8% agarose gel (upper panel) and visualised with EtBr post-staining in a Gel Doc (Bio-Rad). The other half was denatured and labelled with T4 PNK and ^32^P-γ-ATP, analysed by 15% denaturing PAGE and visualised by phosphorimaging in a Typhoon FLA 9500. (-) indicates no enzyme. The arrow indicates products of expected sizes.

However, we were still surprised by the level of non-coordinated cleavage that Yen1 was able to promote, as evidenced by the presence of the relaxed form of the plasmid in these experiments. Given our previous observations of Yen1 arm-chopping activity, we wondered whether this activity could also act on a cruciform-bearing plasmid. If so, it would imply a coordinated pair on incisions, but in an asymmetrical manner, that could lead to the formation of a relaxed, gapped circle (Figure 6A, central row), a product that would be electrophoretically indistinguishable from a relaxed, nicked circle in an agarose gel. To elucidate if Yen1 gives rise to nicked circles (uncoordinated cleavage) or gapped circles (coordinated cleavage), we designed two different experiments. First, we argued that if the relaxed circles were nicked molecules, they could be religated; if they contained ssDNA gaps, they should not religate. As shown in Figure 6C, nicked circles produced with Nt.BspQI are readily ligated, and migrate as a supercoiled DNA band, while only a minor fraction of the relaxed circle can religate in the reactions with Yen1 (Figure 6C). This suggests that the main circular product generated by Yen1 could be a gapped molecule. However, since such gapped circles might arise from nicks extended into gaps by Yen1 itself through its exonuclease activity, a second experiment was designed to specifically detect the DNA fragment that would be released only if Yen1 catalyzed an arm-chopping reaction on one of the hairpins of the cruciform structure (Figure 6D), by labelling all the reaction products and analysing them in a denaturing gel. For the experiment, the cruciform-based plasmid was incubated with a variety of enzymes: 3 different versions of Yen1 (Yen1^WT-λ^, Yen1^ON-λ^ and Yen1^ND^), GEN1 as a control for a linear product, Nt.BspQI as a control for a nicked product, and EcoRI, which is commonly used to test the extrusion level of the plasmid. When these reactions were analysed in a denaturing gel, we could only detect bands consistent with the expected size of the released hairpin (66 nt) in those reactions containing active Yen1, but not in any of the other controls (Figure 6E). All these results strongly suggest that a significant fraction of the relaxed, circular molecules arise from an arm-chopping mechanism. Therefore, Yen1 seems to mainly promote the coordinated incision of plasmid-borne cruciform structures, albeit occasionally in an asymmetrical manner.

### The assembly of a Yen1 dimer on the HJ accelerates the rate of the first incision during HJ resolution

Our results with the plasmid-borne cruciform indicate that HJ processing by Yen1 proceeds by the introduction of coordinated incisions within the lifetime of a nuclease dimer-HJ complex. For the dimeric prokaryotic and mitochondrial resolvases such coordination is ensured by the acceleration of the cleavage of the second strand after the first incision (92). However, eukaryotic Rad2/XPG family HJ resolvases are monomeric in solution and dimerization happens on the HJ (38, 48, 49, 51) (Supp. Fig. 4). It has been shown for human GEN1 that dimerization on the HJ substrate stimulates the first incision of the junction by one of the monomers, which is the rate-limiting step for HJ resolution (38), while experiments with CtGEN1 have confirmed that cleavage of the second strand is accelerated after the first incision (51). To test if the formation of the dimer on a HJ could also favour the initial incision by Yen1, we created a modified X0 HJ with an SP bond at the expected cleavage position of Yen1 in oligo X0-3, between nucleotides 31 and 32 (Figure 7A). When incubated with Yen1 or GEN1, this X0-SP cannot be resolved as efficiently as a normal X0 junction (Figure 7B), mostly due to a drastic decrease in the cleavage of strand X0-3 (Figure 7C), as previously shown for GEN1 (38). Then, we titrated a binding proficient, but catalytically inactive version of Yen1 (Yen1^ND^) in reactions with limiting amounts of active Yen1 to see how an increase of total dimers in the reaction would affect the initial incision. In agreement with published results for GEN1, the nicking of strand X0-1 was stimulated by addition of Yen1^ND^, up to a 1:4 WT:ND molar ratio (Figure 7D and E). As expected, further titration resulted in an inhibitory effect, likely due to the sequestration of HJs by inactive dimers. These results indicate that formation of a Yen1 dimer on the HJ favours the rate-limiting, initial incision.

**Figure 7.**
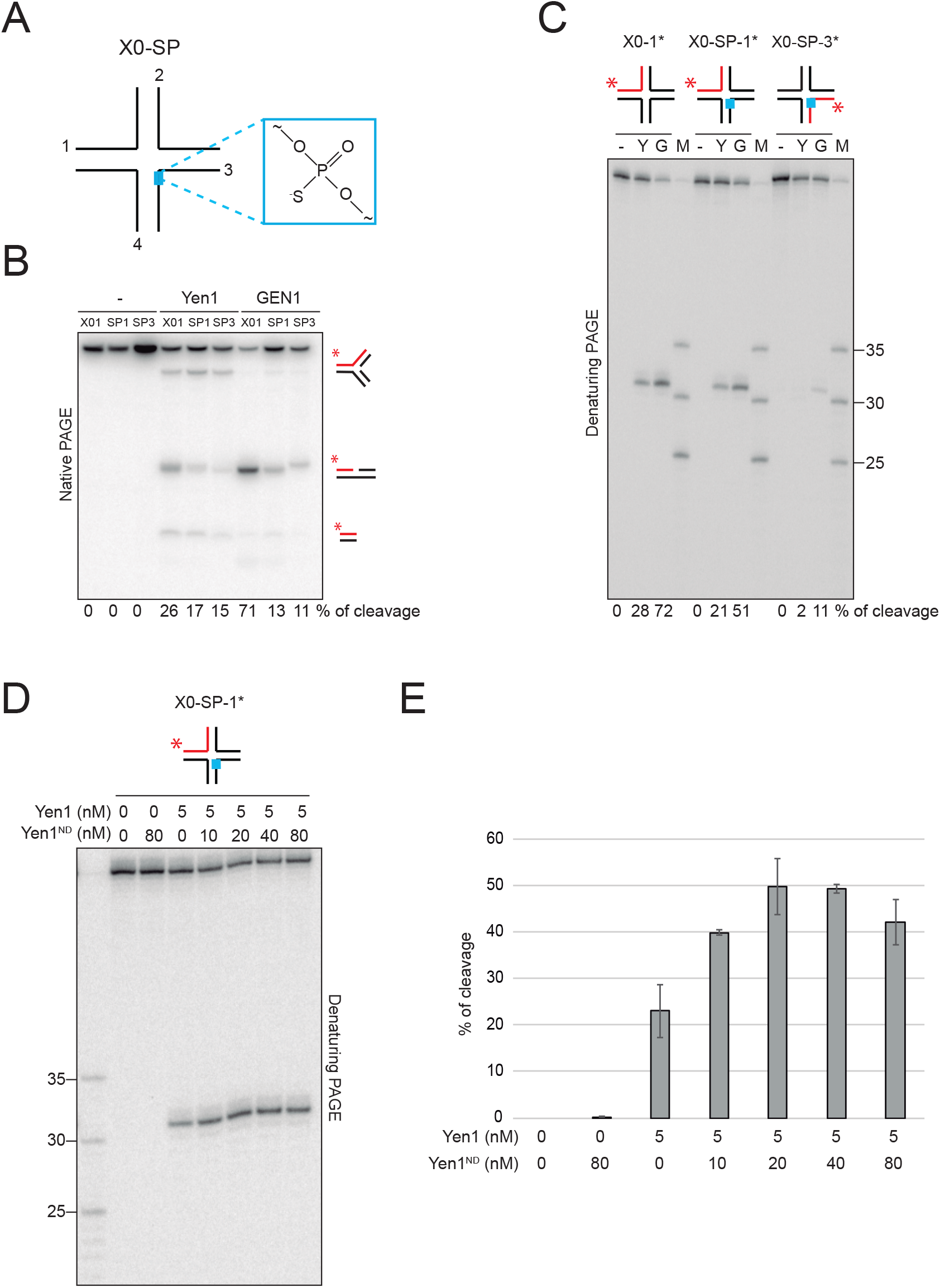
Yen1 dimerization on the substrate favours its first incision. (A) Schematic representation of a modified X0 with a phoshorothioate linkage (in blue) between 31-32 nt of oligo 3 (X0-SP). (B) 5 nM unmodified X0 5’-^32^P-end labelled in oligo 1 (X01) and X0-SP 5’-^32^P-end labelled in oligonucleotide 1 (SP1) or 3 (SP3) were incubated with 5 nM Yen1^λ^ for 5 min at 30°C or 5 nM GEN1^1-527^ for 2.5 min at 37°C. (-) indicates control reaction with no enzyme. Products were analysed by 10% native PAGE. Possible reaction products are indicated on the right. Cleavage was quantified as shown. (C) Same reactions as in (B) were analysed by 10% denaturing PAGE. DNA was labelled in the indicated strands (red). 5’-^32^P-end-labeled oligos of defined length (25, 30 and 35 nt) were used as markers (M). (D) X0-SP (5 nM) 5’-^32^P-end labelled in oligo 1 was incubated with 5 nM Yen1^λ^ and the indicated concentrations of Yen1^ND^ for 5 min at 30°C. Products were analysed by 10% denaturing PAGE. (E) Quantification of the reaction shown in (D). Data represented as mean values ± SD (n = 3).

### Yen1^ON^ displays the same cleavage specificity as Yen1

Yen1^ON^ is a mutant version of Yen1 where all nine serines in CDK consensus sites (Figure 1A) have been substituted for alanines, rendering the protein constitutively active and nuclear throughout the cell cycle (60). This mutant serves as a powerful tool to explore the consequences of its premature activation without the need of interfering with the cell cycle progression machinery (66, 67, 71–78). While we have previously shown that the range of substrates that Yen1^ON^ can process overlaps entirely with that of Yen1 (67, 68), we did not know whether the specificity of the incisions within each substrate remained unaltered, so we mapped the cleavage sites produced by Yen1^ON^ on the same set of substrates (Supp. Fig. 5A). We did not observe any difference on the position of the sites cleaved by Yen1^ON^ with respect to those cleaved by the wild-type enzyme, confirming that despite its lack of CDK-dependent regulation, Yen1^ON^ retains the same cleavage specificity, thus representing a good proxy for the status of maximum Yen1 activation. However, when comparing the kinetics of *in vitro*-dephosphorylated wild-type Yen1 with those of Yen1^ON^, we noticed that the cleavage rate of the former was always lower (Supp. Fig. 5B and C). In this sense, previous phosphomapping analyses of Yen1 revealed that there are at least 20 S/T residues at non-CDK consensus sites phosphorylated *in vivo* (67, 68, 93). Since the Yen1^ON^ preps we employed were not *in vitro*-dephosphorylated, we wanted to assess if phosphorylation at those non-CDK consensus sites could also be contributing to the regulation of Yen1 activity. For this, we dephosphorylated Yen1^ON^ by treatment with lambda-phosphatase and compared its ability to process different substrates with respect to the mock-treated protein (Supp. Fig. 5D and E). Both versions displayed similar cleavage kinetics, indicating that phosphorylation at CDK sites represent the major regulatory switch for Yen1 nuclease activity, with none or minor contribution from other phosphoresidues. Since this result did not explain the differences in activity between dephosphorylated Yen1 and Yen1^ON^, we tested if some level of residual phosphorylation could still be detected for these proteins after lambda-phosphatase treatment using Phos-Tag containing SDS-PAGE gels that specifically reduce the mobility of phosphoproteins. Indeed, we observed that while the phosphatase treatment apparently abolished all phosphorylation on Yen1^ON^, a detectable shift was still present for wild-type Yen1 (Supp. Fig. 5F). Therefore, the more likely explanation for the difference in activity between dephosphorylated wild-type Yen1 and Yen1^ON^ is due to the inability of lambda phosphatase treatment to fully remove all phosphorylation from relevant serines at Yen1 CDK consensus sites and thus preventing full activation of the enzyme.

### Expression in yeast of constitutively nuclear human GEN1^nuc^ does not fully recapitulate the genotoxicity of misregulated Yen1

The ability of both Yen1 and GEN1 to target branched DNA molecules is tightly controlled throughout the cell cycle. To understand the importance of such regulation, two mutants that can access and cleave their potential targets at any cell cycle stage - Yen1^ON^ and GEN1^nuc^ – were expressed in budding yeast and human cells, respectively (50, 67). Interestingly, while expression of both these mutants lead to an increase of COs, only Yen1^ON^ causes hypersensitivity to genotoxic stress. To figure out whether this higher toxicity of Yen1^ON^ expression in yeast cells depends on some specific feature of the yeast chromatin biology or on the biochemical differences between Yen1 and GEN1, we decided to set up an *in vivo* experiment to compare the effects of the expression of these mutants in the same organism. For this, we created a series of *S. cerevisiae* strains harboring different cassettes to express Yen1, Yen1^ON^, GEN1, GEN1^nuc^ and the GEN1 truncated versions GEN1^1-527^ and GEN1^1-527nuc^, all under the control of either the inducible *GAL1* or the constitutive *GPD1* promoters in both wild-type and resolution-deficient (*mus81Δ yen1Δ*) backgrounds. To ensure that any observed toxicity for Yen1^ON^ was not merely due to a dose or localization effect, we aimed for a set of strains where GEN1 variants displayed constitutive nuclear localization as well as similar or higher protein expression levels and similar or higher biochemical activity in whole-cell extracts when compared to strains carrying Yen1^ON^. Among all combinations tested, these premises could only be fulfilled in media containing galactose when Yen1 and Yen1^ON^ were expressed under the control of the *GPD1* promoter, and GEN1^1-527^ and GEN1^1-527nuc^ were expressed under the control of the *GAL1* promoter (Figure 8A-C). Under these conditions, we compared the effect of Yen1^ON^ and GEN1^1-527nuc^ in increasing hypersensitivity to the genotoxic agent methyl methanesulfonate (MMS) in a wild-type background (Figure 8D). To verify the functionality of the enzymes, we also performed this experiment in the resolution-deficient *mus81Δ yen1Δ* double mutants (Figure 8E), where premature activation of Yen1 provides increased resistance to MMS (67). In the WT strain, expression of Yen1^ON^ conferred strong sensitivity at the lower dose of MMS tested (Fig. 8D), whereas expression of GEN1^1-527nuc^ induced a milder sensitivity, despite showing comparable expression levels and a more potent nuclease activity on extracts (Fig. 8A and B). However, when we analysed the survival of the *mus81Δ yen1Δ* double mutants to the MMS treatment, we observed that Yen1^ON^ was also more efficient in providing resistance to MMS than GEN1^1-527nuc^ (Figure 8E). Therefore, while Yen1^ON^ expression is more toxic than that of GEN1^1-527nuc^, despite its lower biochemical activity in whole cell extracts, we cannot conclusively affirm that this toxicity derives from specific differences in their biochemical properties, as GEN1^1-527nuc^ does not seem comparably functional in the processing of HR intermediates when expressed in yeast.

**Figure 8.**
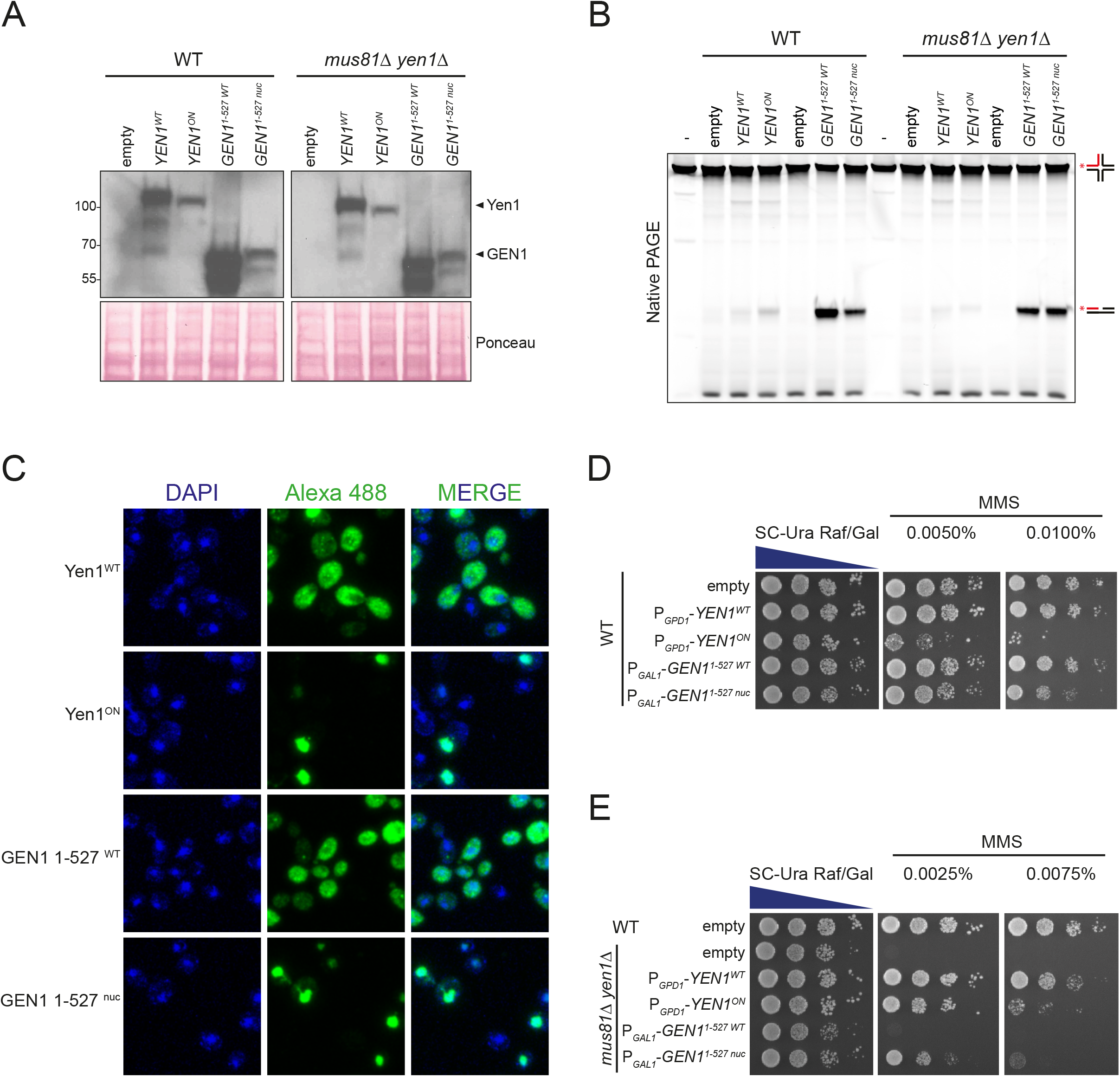
Expression of deregulated Yen1 and GEN1 mutants in *S. cerevisiae*. (A) Western-blot analysis of protein expression levels in soluble extracts from WT or *mus81Δ yen1Δ* strains transformed with the plasmids p416GPD1-YEN1^WT/ON^-V5-6xHis or pEXP52-GEN1^1-527^ ^WT/NUC^-V5-6xHis growing in SC-URA containing 1% raffinose and 2% galactose. Proteins were detected with an α-V5 antibody and Ponceau staining was employed as a loading control. (B) Resolution assay employing soluble extracts from (A). 2 µg total protein was incubated with 50 nM of IRDye-700 fluorescent X0-1* for 1 h at 30°C. Products were analysed in 10% native PAGE and visualised in an Odyssey (LI-COR) infrared scanner. (C) Analysis of the subcellular localization of Yen1 and GEN1 variants by immunofluorescence employing α-V5 (1:200) as primary antibody and α-mouse Alexa 488 (1:200) as secondary antibody. DNA was stained with DAPI. (D) Wild-type strains carrying the indicated constructs were plated on media with increasing amounts of MMS, grown at 30°C for 72 h and photographed. (E) As in (D), but employing *mus81Δ yen1Δ* strains.

## DISCUSSION

The original identification of Yen1 as a canonical HJ resolvase in *S. cerevisiae* relied on experiments carried out employing whole cell-extracts and protein immobilized on agarose beads (10). Since then, much of what we know about Yen1 properties has been extrapolated from other orthologs in the subclass IV of the Rad2/XPG family that were successfully purified before. Despite the subsequent purification of functional Yen1 to homogeneity, only a basic study of its mechanism of activation was described (67). Here, we presented the first exhaustive biochemical characterization of Yen1, focused on the evaluation of its properties compared to the well-studied human canonical HJ resolvase GEN1.

### The high- and low-activity versions of Yen1 display identical incision patterns

We had previously shown that the phosphorylation status of Yen1 does not affect the range of substrates it can target (67), which are similar to those described for GEN1 (10, 38, 54). Here, we have additionally confirmed that these changes in phosphorylation do not alter the position of the incisions that Yen1 produces on a wide range of substrates (Figure 1, Supp. Fig. 1 and 2). In a similar way, the constitutively active mutant Yen1^ON^ incises all the tested substrates at identical positions when compared to the wild-type Yen1 (Supp. Fig. 5). Therefore, this work further substantiates that the activation of Yen1 by dephosphorylation does not qualitatively change how the enzyme processes these oligonucleotide-based substrates.

### Yen1 may present a particular interaction mode with HJs and other substrates

In addition to some of the distinct incisions in various substrates that we have observed, a particularly striking difference between Yen1 and GEN1 is their opposed preference for the crossing and continuous strands, respectively, of the J3 HJ (Figure 4). Despite its closer phylogenetic relation to Yen1, a preference for the continuous strands has also been observed in the fungal CtGEN1 (51). This suggests that Yen1 might bind or remodel the junction in different ways compared to GEN1, CtGEN1 and, potentially, other orthologs. While structural details are not available for Yen1, we have identified a *Saccharomyces* genus-specific, disordered region in the XPG-N domain in Yen1, whose deletion renders the protein inactive (our unpublished data). This suggests that this region, which is located near essential catalytic residues like D41, could be to some extent involved in the positioning of the catalytic residues and hence modify the way Yen1 specifically interacts with the HJ with respect to other orthologs from other genera.

We would also like to note that we have detected unanticipated cleavage products by human GEN1 on oligo X0-1 in substrates like the 3’-flap, splayed arm or ssDNA (Supp. Fig. 2). These products can be observed in GEN1 preparations from other groups (49). Since this activity can be observed on X0-1 as ssDNA, but not on ssDNA poly-T oligos, we think that this oligo may form a stable secondary structure that can be recognized and targeted by GEN1, but not Yen1, revealing another difference between the two enzymes.

### Yen1 non-canonical activities and the paradigm of canonical HJ resolution

The paradigm of canonical resolution states that the symmetrical, coordinated incisions across one of the axis of the junction must occur within the lifetime of the enzyme-substrate complex, yielding nicked DNA molecules that can be sealed without the need for further processing (41, 84). This work demonstrates that Yen1 can be considered a canonical resolvase since i) it can create symmetrical cuts on a HJ (Figure 1), ii) the incisions occur mainly in a coordinated manner (Figure 6) and iii) its resolution products, in a first instance, could be ligated (Figure 5). However, the existence of the novel arm-chopping and nick-exonuclease activities plays a confounding role in the interpretation of biochemical assays traditionally employed to assess the coordination of incisions and the ligation of resolution products, respectively.

In this sense, our results regarding cleavage coordination with the cruciform extruding plasmid pIR9 (48) led us to propose an extension to the usual interpretation for this type of experiments, where an undetermined, but likely major fraction of the relaxed circular species produced by Yen1 corresponds to gapped, rather than nicked circles, resulting from arm-chopping activity (Figure 6 and Supp. Fig. 3B). In this way, Yen1 would process this cruciform mainly in a coordinated, albeit not always symmetrical manner. This coordination would be consistent with the idea that Yen1, despite being a monomer in solution (Supp. Fig. 4), needs to dimerize on the substrate in order to accelerate the rate of the first incision (Figure 7C-D), the limiting step of the reaction, as previously proposed for GEN1 in humans, *D. melanogaster, C. thermophilum* and other eukaryotic resolvases (38, 40, 46, 47, 49, 54, 94). With respect to the nick-exonuclease activity, it is interesting to note that it has also been detected for GEN1, albeit it does not seem robust enough to prevent the majority of the resolution products from being religated (Figure 5E and F). Importantly, we have never observed exonuclease activity of Yen1 on dsDNA or ssDNA substrates (Supp. Fig. 2), reinforcing the notion that this activity is specific of nicked dsDNA molecules. At this point we cannot categorically rule out that such activity stems from partial DNA breathing that creates pseudo-5’flaps at the nicks that are processed endonucleolytically. However, if this was the case, we would expect that GEN1, being generally more active on 5’-flaps than Yen1, should also display a stronger activity on the nicked dsDNA, which is not the case.

Altogether, our results support that Yen1 has a canonical resolvase behaviour *in vitro*, despite the presence of two additional activities that may blur such classification.

### Could the detrimental effects of Yen1 premature activation be due to non-canonical activities operating *in vivo*?

Yen1 and GEN1 ability to access and process their targets in chromatin is strictly regulated throughout the cell cycle (62, 63, 67–69). However, the misregulation of Yen1 seems to elicit a more detrimental effect than misregulation of GEN1, especially in terms of DNA damage sensitivity (50, 67). Such effect could be explained either by Yen1 deregulation having more severe consequences due to its specific activities or by yeast system being generally more sensitive to misregulation of any nuclease. Our efforts to try to distinguish between these two possibilities have not allowed us to reach conclusive results, if interpreted conservatively (Figure 8). The expression in yeast of a deregulated version of GEN1, GEN1^1-527nuc^, produces a milder phenotype than Yen1^ON^ both in the increase of sensitivity to MMS in wild-type cells and in increased resistance to MMS in *mus81Δ yen1Δ* double mutants. These results would suggest that GEN1^1-527nuc^ does not have a similar capacity to target branched DNA molecules to Yen1^ON^ in yeast, regardless of their physiological or pathological nature for the cell. Notwithstanding, it has been shown that expression of GEN1 can process repair intermediates and restore sporulation in *S. pombe mus81Δ* mutants (95), and the difference in the detrimental effect between Yen1^ON^ and GEN1^1-527nuc^ expression in wild-type cells seems overtly higher than in their protective effect in *mus81Δ yen1Δ* strains (Figures 8D and 8E). Therefore, we consider that, to some extent, Yen1 may present specific properties that lead to increased toxicity when prematurely activated compared to GEN1. However, whether this potential toxicity of Yen1 is due to the novel activities described in this work, or if they actually manifest *in vivo*, remains unclear. The nicked dsDNA-specific 5’-3’exonuclease activity described in this paper has been previously described for other members of the Rad2/XPG family, like class II human FEN1 (96, 97) and class III EXO1 (98), and even in the class IV Yen1/GEN1 ortholog DmGEN (43). Additionally, even if not specifically tested, the low ligation efficiency observed for the resolution products of OsGEN-L, AtGEN1 and AtSEND1 (48, 56), could arguably be due to the presence of a similar activity. With respect to Yen1 ability to excise one arm from a nicked HJ (*Ref-I* activity (48)) or an intact HJ (arm-chopping) (Supp. Fig. 3), at least one of these activities has also been described or can be observed in the experiments with the plant, fly and mammalian orthologs (48, 49, 56) (Figure 2 and Supp. Fig.1), and hence seem rather widespread among the subclass IV of the Rad2/XPG family of nucleases. Despite this apparent conservation, it is not straightforward to envisage a possible biological role for an activity like the arm-chopping in the context of disengaging two DNA molecules connected by HJs, as it would inevitably lead to the generation of a DSB (Supp. Fig. 3). Alternatively, it has been proposed that replication fork reversal, which leads to the formation of a HJ-like structure, may help stabilize stalled replisomes, sometimes leading to situations of transient over-replication (99). Then, template unwinding followed by flap processing or end-resection of the regressed arm would be required for its removal and replication termination. In this context, if such intermediates persisted until late stages of mitosis, the endonucleolytic processing of the regressed arm by arm-chopping could hypothetically provide an alternative way for its elimination. In the future, it would be interesting to take advantage of structural data to explore possible separation-of-function mutants that lack some of these non-canonical activities. With such tools in hand, we could ask if they actually play any functional role *in vivo*, like the processing of particularly adamant recombination or replication intermediates, or simply represent collateral consequences of the evolutionary adaptation of a monomeric SSE to perform a reaction that requires its dimerization and coordinated cleavage on a Holliday junction.

## SUPPLEMENTARY DATA

Supplementary data contains Supplementary Tables 1-4 and 5 Supplementary Figures.

### ACKNOWLEDGEMENTS

The authors would like to thank Gary Chan and Stephen C. West for various yeast strains, plasmids and proteins; Daniela Kobbe for the pIR9 plasmid; Joao Matos and Tomás Lama Díaz for constructive discussions and for critical reading of the manuscript and Marta Picado for technical assistance with fluorescence microscopy.

## SUPPLEMENTARY DATA

Supplementary data contains Supplementary Figures 1-5 and Supplementary Tables 1-4

## Supplementary Figure legends

**Supplementary Figure 1.**
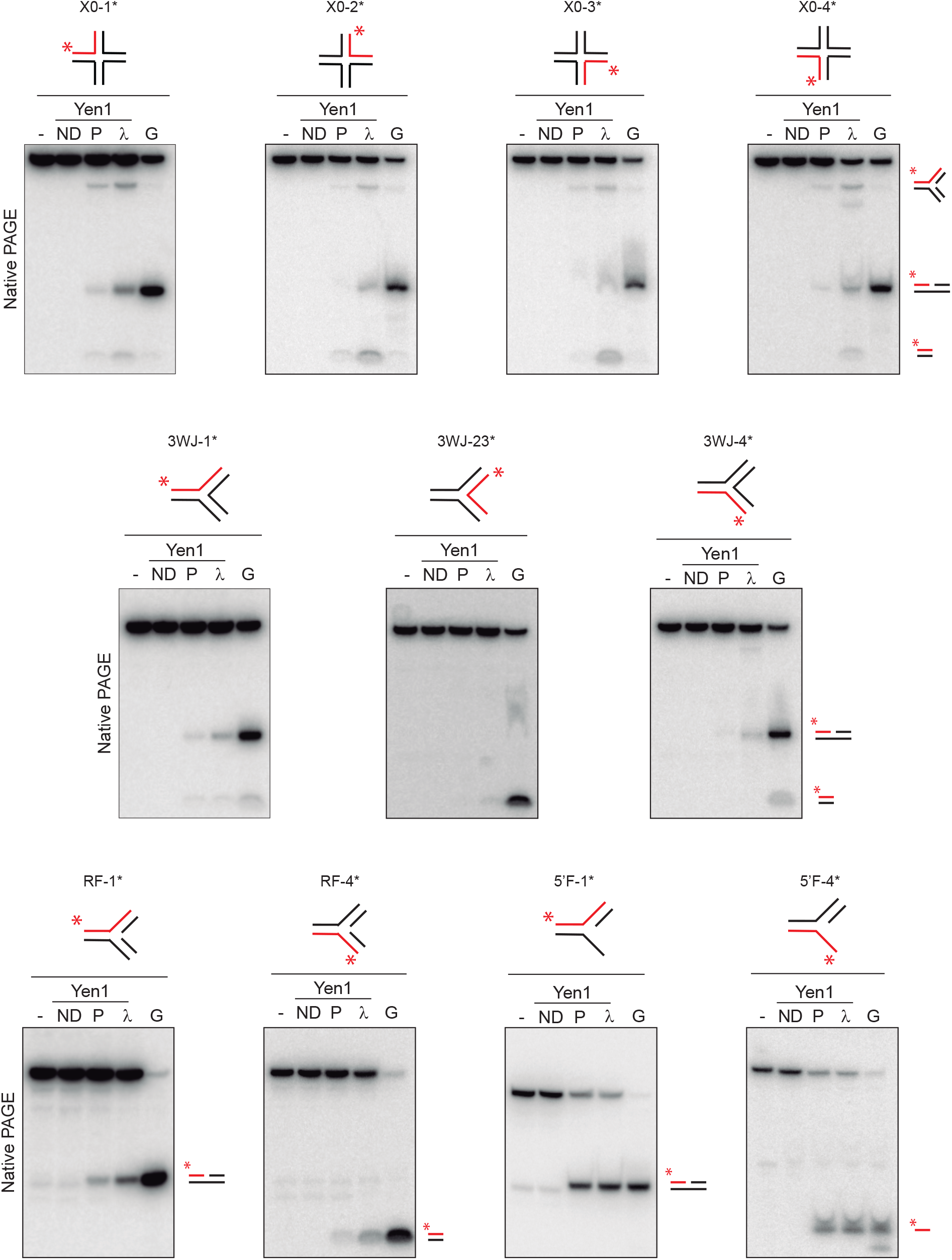
Range of substrates cleaved by Yen1. The same reactions shown in Figure 1C-F were analysed in 10% native PAGE and phosphorimaging. Reaction products are depicted on the right. All labels as in Figure 1.

**Supplementary Figure 2.**
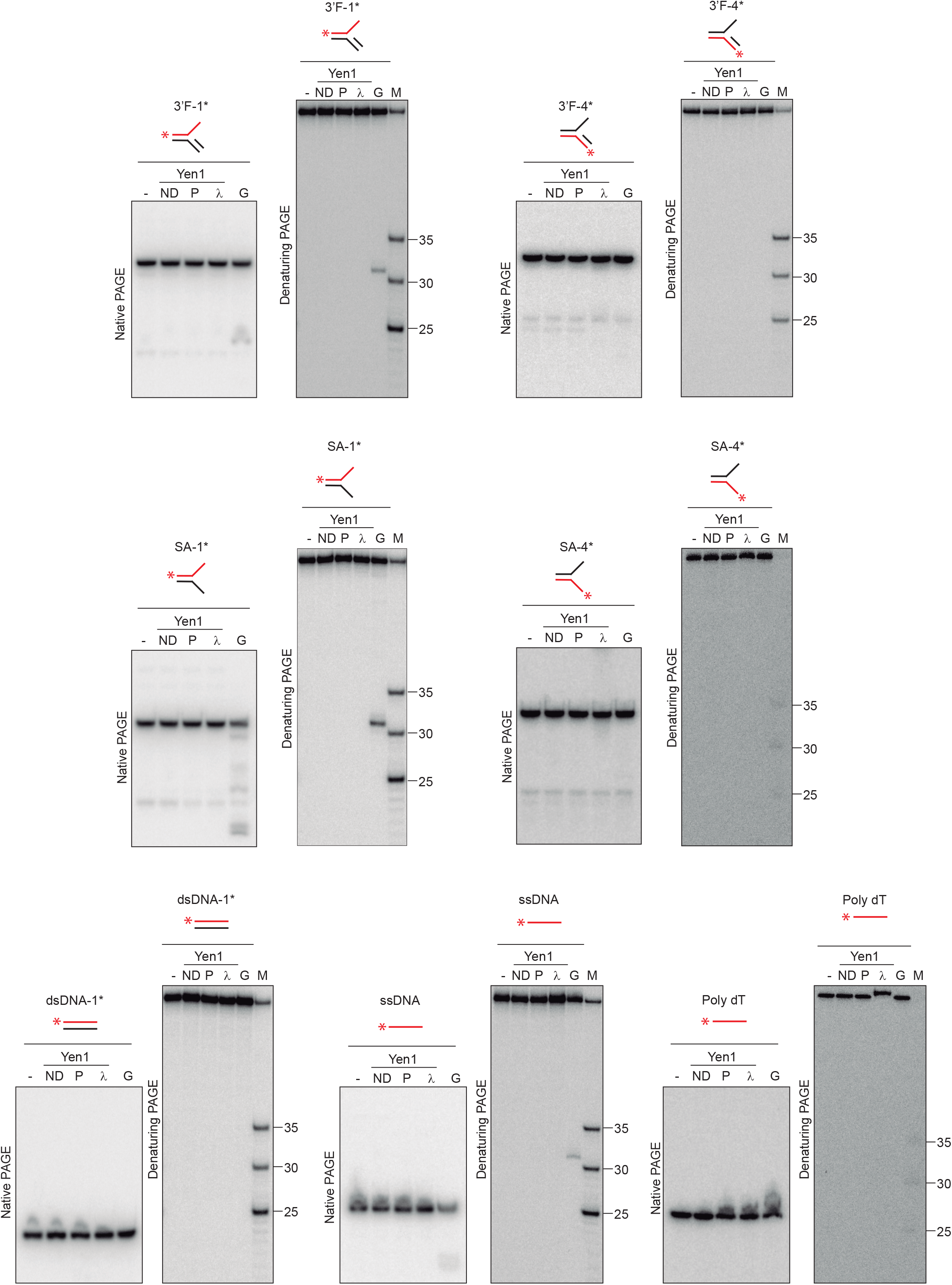
Substrates not processed by Yen1. Reactions as those described for Figure 1C-F and Supp. Fig.1 were carried out employing the following substrates: 3’-flap (3’F), splayed arm (SA), dsDNA, ssDNA and poly-dT. All substrates were 5’-^32^P-end labelled at the indicated strand (red) and incubated with Yen1^ND^, Yen1^P^, Yen1^λ^ and GEN1^1-527^ (5 nM) as described. Reaction products were analysed by 10% denaturing or native PAGE and phosphorimaging. All labels as in Figure 1 and Supp. Fig. 1.

**Supplementary Figure 3.**
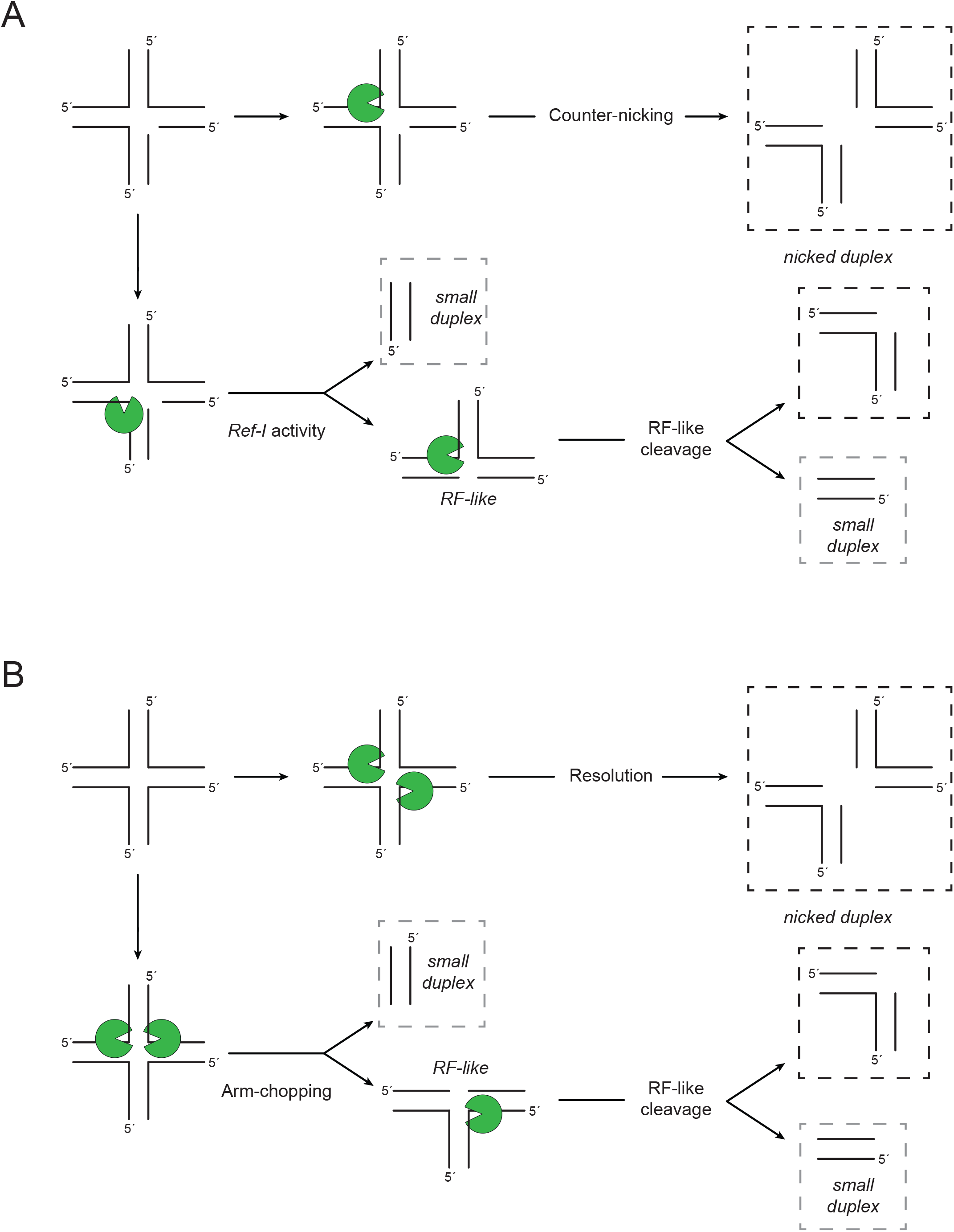
Graphic representation of the different modes for the processing of nicked and fully ligated HJs. (A) Nicked HJs can be counter-nicked, presumably by a Yen1 monomer, in the strand opposite the nicked strand, releasing two nicked dsDNA molecules (top row). Alternatively, the incision may occur on a strand that anneals with the nicked strand (*Ref-I* activity), releasing a short dsDNA duplex and a replication fork-like (RF-like) structure, which can be further cleaved into a nicked dsDNA molecule and a second short dsDNA duplex (bottom row). (B) Intact HJs may be canonically resolved through coordinated incisions on opposite strands by a Yen1 dimer to release two nicked dsDNA molecules (top row). If coordinated cleavage occurs by coordinated incisions on annealing strands (arm-chopping), a short dsDNA duplex and a replication fork-like (RF-like) structure are released similarly to *Ref-I* activity. Further processing of the RF-like would also generate a nicked dsDNA molecule and a second short dsDNA duplex (bottom row).

**Supplementary Figure 4.**
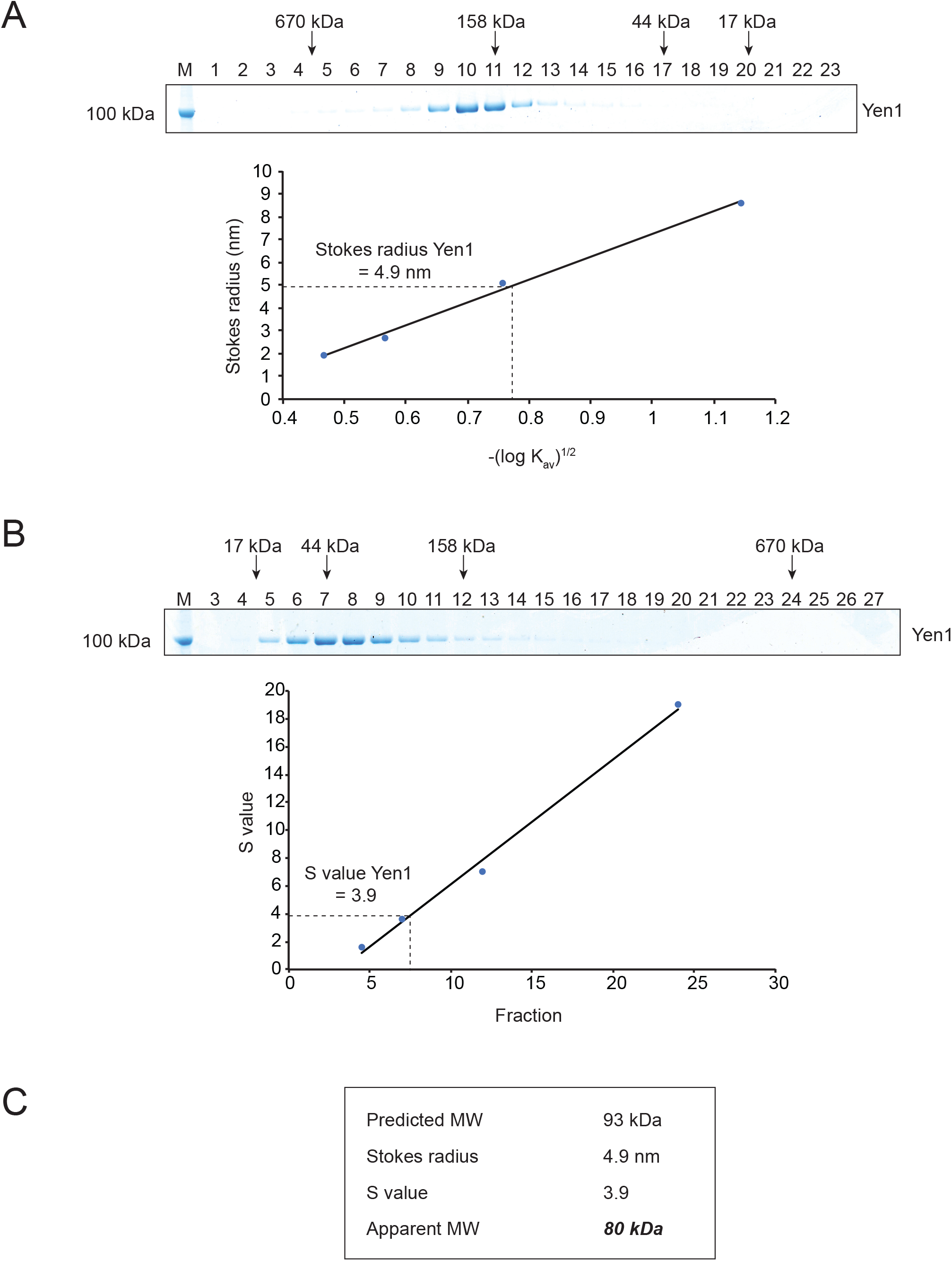
Yen1 is a monomer in solution. (A) Determination of Yen1 Stokes radius. Fractions from size exclusion chromatography of 20 *µ*g Yen1^λ^ were analysed by SDS-PAGE and Coomassie staining. The peak fraction number where filtration standards (thyroglobulin, γ-globulin, ovalbumin and myoglobin; BioRad) eluted is indicated by arrows. The observed K_av_ values and known Stokes radii of the standards were employed to calculate the Stokes radius for Yen1. (B) Determination of Yen1 sedimentation coefficient. Fractions collected after ultracentrifugation of 25 *µ*g Yen1^λ^ through glycerol gradients were analysed as in (A) and the peak fraction for each MW standard plotted to estimate Yen1 S-value. (C). Estimation of the apparent MW for Yen1 from the observed hydrodynamic values employing the Siegel and Monty equation (1). The calculated MW for FTH-tagged Yen1 is 93 Kda.

**Supplementary Figure 5.**
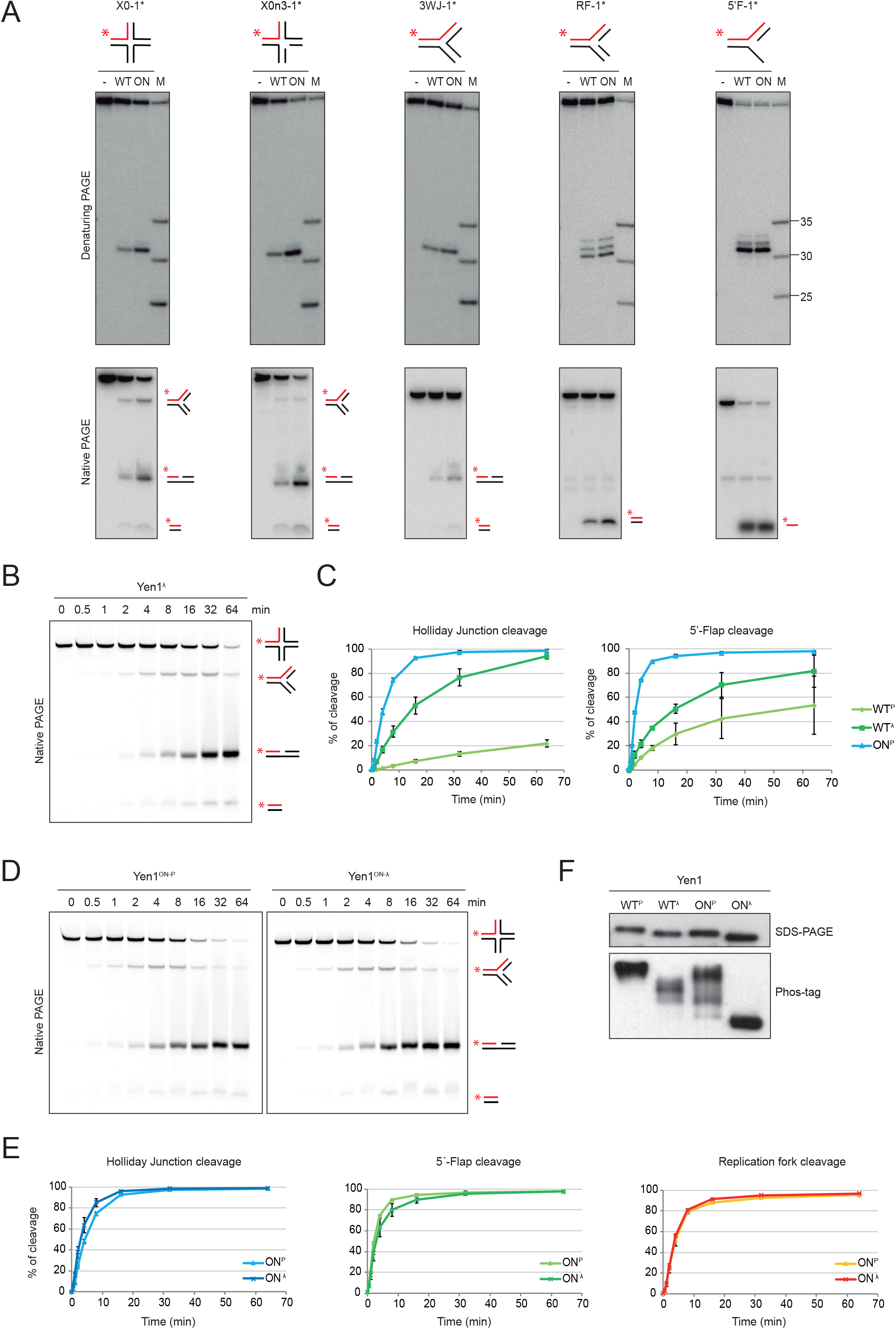
Comparison of the activity of Yen1 and Yen1^ON^. (A) Reactions as described in Figure 1C-F and Supp. Fig 1 were carried out with 20 nM substrate and 5 nM Yen1^λ^ (WT) or Yen1^ON^ (ON) for 10 min at 30°C. Reaction products were analysed by 10% denaturing (top) or native (bottom) PAGE and phosphorimaging. All labels as in Figure 1 and Supp. Fig. 1. (B-C) Time-course analysis of Yen1^P^, Yen1^λ^ and Yen1^ON-P^ activity on HJ and 5’-flap substrates. 50 nM of an IRDye-800 fluorescent X0 HJ or 10 nM 5’-flap were incubated with 40 nM of Yen1^P^, Yen1^λ^ or Yen1^ON-P^ for different time points (0, 0.5, 1, 2, 4, 8, 16, 32, 64 min). Products were analysed in 10% native-PAGE and visualised by phosphorimaging as stated above. Figure only shows a representative image of the experiments. (C) Quantification of the gels described in (B). Data represents mean ± SD from two independent experiments. (D) Same as in (B), but the image representing the assay with Yen1^ON-P^ and Yen1^ON-λ^. (E) Quantification of gels in (D). Data represents mean ± SD from two independent experiments. (F) Comparison of the phosphorylation status of different Yen1 versions. 1 µg of FTH-tagged Yen1^P^, Yen1^λ^, Yen1^ON-P^ or Yen1^ON-λ^ was loaded in two independent SDS-PAGE gels, one of them containing Phos-Tag to specifically reduce the mobility of phosphorylated proteins. Proteins were detected using α-FLAG-HRP antibody.

## Supplementary Tables

**Supplementary Table 1.**
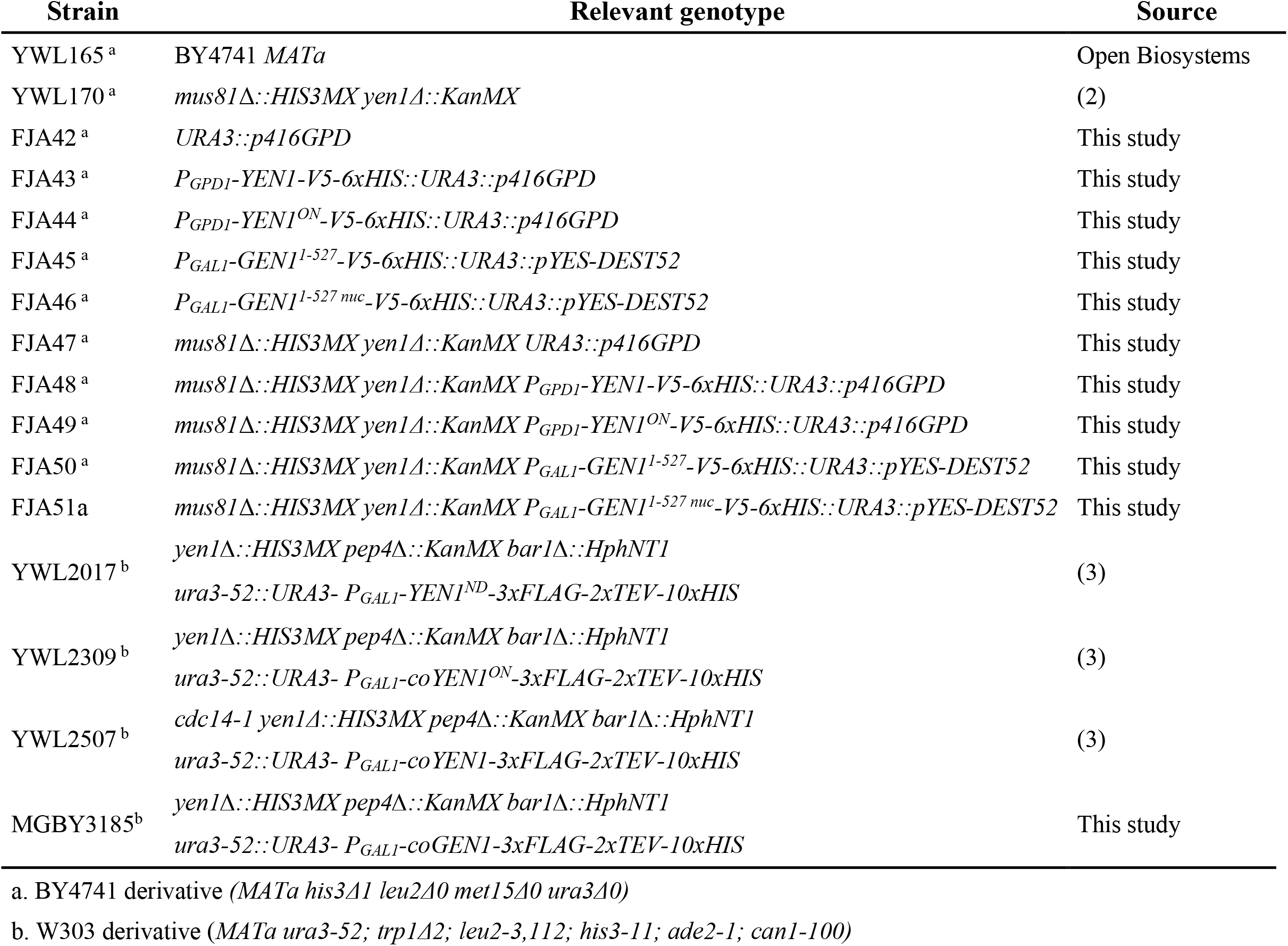
*S. cerevisiae* strains used in this study.

**Supplementary Table 2.** Plasmids used in this study.

| Plasmid | Source |
| --- | --- |
| pIR9 | (4) |
| p416GPD1 | (5) |
| p416ADH1-YEN1<sup>WT</sup>-V5-6xHis | (2) |
| p416ADH1-YEN1<sup>ON</sup>-V5-6xHis | This work |
| p416GPD1-YEN1<sup>WT</sup>-V5-6xHis | This work |
| p416GPD1-YEN1<sup>ON</sup>-V5-6xHis | This work |
| pYES-DEST52 | Invitrogen |
| pAG306GAL-ccdB | (6) |
| pENTR221-GEN1<sup>WT</sup>-FTH-STOP | (7) |
| pENTR221-GEN1<sup>nuc</sup>-FTH-STOP | This work |
| pENTR221-GEN1<sup>WT</sup>-GO | This work |
| pENTR221-GEN1<sup>nuc</sup>-GO | This work |
| pYES-EXP52-GEN1<sup>WT</sup>-V5-6xHis | This work |
| pYES-EXP52-GEN1<sup>nuc</sup>-V5-6xHis | This work |
| pYES-EXP52-GEN1<sup>1-527 WT</sup>-V5-6xHis | This work |
| pYES-EXP52-GEN1<sup>1-527 nuc</sup>-V5-6xHis | This work |
| pAG306GAL-GEN1<sup>WT</sup>-FTH | This work |

**Supplementary Table 3.**
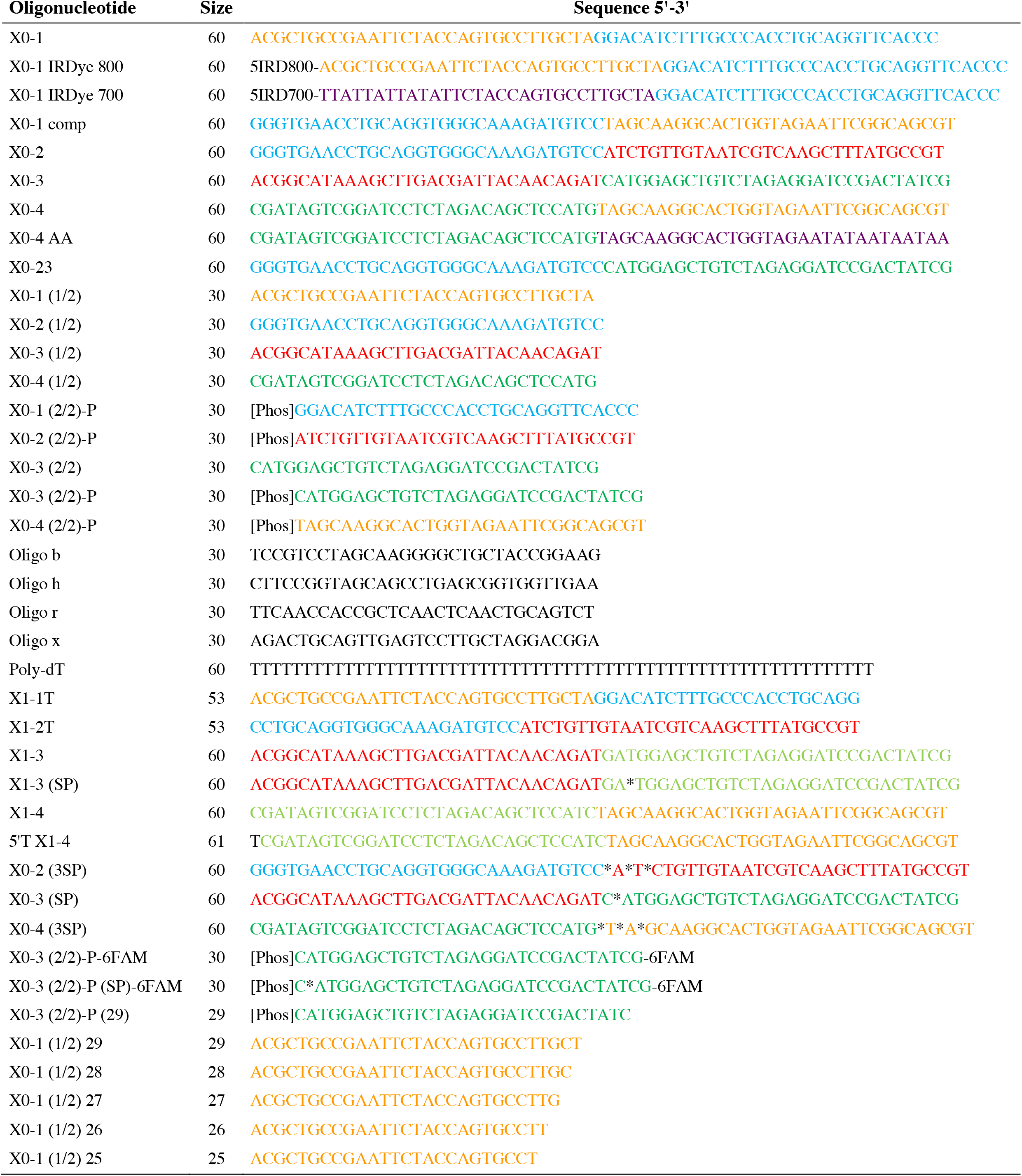
Sequence of the oligonucleotides used for synthetic DNA substrates.

**Supplementary Table 4.**
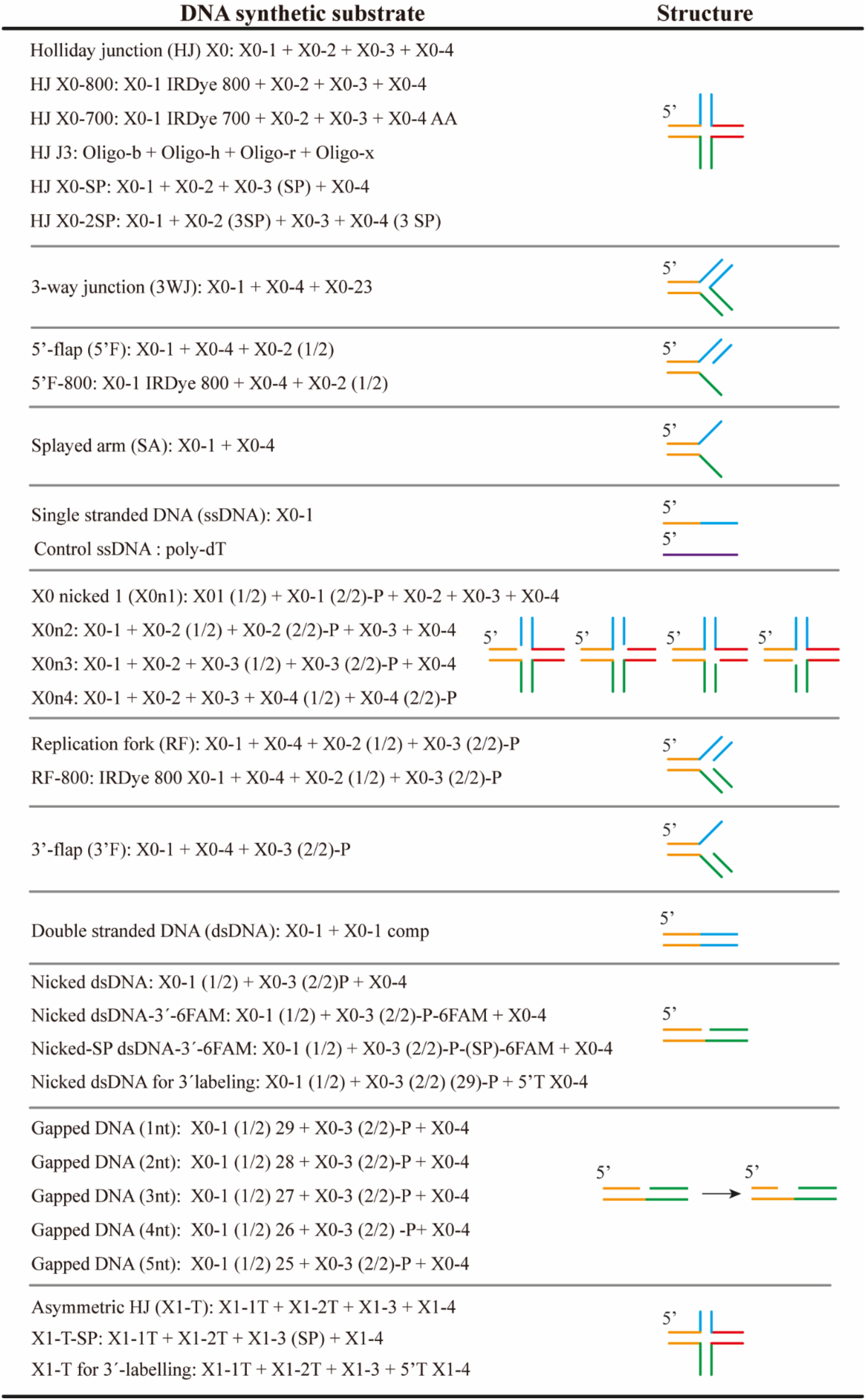
Oligonucleotide combination to generate the different DNA substrates.

| DNA synthetic substrate | Structure |
| --- | --- |
| Holliday junction (HJ) X0: X0-1 + X0-2 + X0-3 + X0-4 |  |
| HJ X0-800: X0-1 IRDye 800 + X0-2 + X0-3 + X0-4 |  |
| HJ X0-700: X0-1 IRDye 700 + X0-2 + X0-3 + X0-4 AA |  |
| HJ J3: Oligo-b + Oligo-h + Oligo-r + Oligo-x |  |
| HJ X0-SP: X0-1 + X0-2 + X0-3 (SP) + X0-4 |  |
| HJ X0-2SP: X0-1 + X0-2 (3SP) + X0-3 + X0-4 (3 SP) |  |
| 3-way junction (3WJ): X0-1 + X0-4 + X0-23 |  |
| 5'-flap (5'F): X0-1 + X0-4 + X0-2 (1/2) |  |
| 5'F-800: X0-1 IRDye 800 + X0-4 + X0-2 (1/2) |  |
| Splayed arm (SA): X0-1 + X0-4 |  |
| Single stranded DNA (ssDNA): X0-1 |  |
| Control ssDNA : poly-dT |  |
| X0 nicked 1 (X0n1): X01 (1/2) + X0-1 (2/2)-P + X0-2 + X0-3 + X0-4 |  |
| X0n2: X0-1 + X0-2 (1/2) + X0-2 (2/2)-P + X0-3 + X0-4 |  |
| X0n3: X0-1 + X0-2 + X0-3 (1/2) + X0-3 (2/2)-P + X0-4 |  |
| X0n4: X0-1 + X0-2 + X0-3 + X0-4 (1/2) + X0-4 (2/2)-P |  |
| Replication fork (RF): X0-1 + X0-4 + X0-2 (1/2) + X0-3 (2/2)-P |  |
| RF-800: IRDye 800 X0-1 + X0-4 + X0-2 (1/2) + X0-3 (2/2)-P |  |
| 3'-flap (3'F): X0-1 + X0-4 + X0-3 (2/2)-P |  |
| Double stranded DNA (dsDNA): X0-1 + X0-1 comp |  |
| Nicked dsDNA: X0-1 (1/2) + X0-3 (2/2)P + X0-4 |  |
| Nicked dsDNA-3'-6FAM: X0-1 (1/2) + X0-3 (2/2)-P-6FAM + X0-4 |  |
| Nicked-SP dsDNA-3'-6FAM: X0-1 (1/2) + X0-3 (2/2)-P-(SP)-6FAM + X0-4 |  |
| Nicked dsDNA for 3' labeling: X0-1 (1/2) + X0-3 (2/2) (29)-P + 5'T X0-4 |  |
| Gapped DNA (1nt): X0-1 (1/2) 29 + X0-3 (2/2)-P + X0-4 |  |
| Gapped DNA (2nt): X0-1 (1/2) 28 + X0-3 (2/2)-P + X0-4 |  |
| Gapped DNA (3nt): X0-1 (1/2) 27 + X0-3 (2/2)-P + X0-4 |  |
| Gapped DNA (4nt): X0-1 (1/2) 26 + X0-3 (2/2) -P+ X0-4 |  |
| Gapped DNA (5nt): X0-1 (1/2) 25 + X0-3 (2/2)-P + X0-4 |  |
| Asymmetric HJ (X1-T): X1-1T + X1-2T + X1-3 + X1-4 |  |
| X1-T-SP: X1-1T + X1-2T + X1-3 (SP) + X1-4 |  |
| X1-T for 3'-labelling: X1-1T + X1-2T + X1-3 + 5'T X1-4 |  |

